# Cryo-EM reveals the membrane binding phenomenon of EspB, a virulence factor of the Mycobacterial Type VII secretion system

**DOI:** 10.1101/2022.09.14.507925

**Authors:** Nayanika Sengupta, Surekha P., Somnath Dutta

## Abstract

*Mycobacterium tuberculosis* utilizes sophisticated machinery called the type VII secretion system to translocate virulence factors across its complex lipid membrane. ESX-1 is one of the essential and well-studied secretion systems which transport various virulence factors, including EspB. EspB, a ~36 kDa secreted substrate, has been implicated to play vital role in protecting the bacteria from hostile environment within the host cell phagosome. It is also involved in bacterial pathogenesis and has been shown to bind phospholipids. Recently, two cryo-EM structures of EspB full-length and the secreted isoforms were resolved. Despite the availability of multiple high-resolution structures of EspB, the physiological relevance and mechanism of virulence of this secreted substrate remains poorly characterized. In this current work, we implemented cryo-EM-based structural studies, including various functional assays, TEM imaging, and biophysical approach to demonstrate the interaction of EspB with lipids and bio-membrane. Our findings also indicated that EspB may play a crucial role in binding to and rupturing host mitochondrial membrane. Through cryo-EM studies we were able to show the possible membrane-binding region of EspB. Our study sheds light on host-pathogen interactions and bacterial pathogenesis mediated by EspB.

## Introduction

*Mycobacterium tuberculosis* (*M. tuberculosis*), a unique Gram-positive bacterium is accounted as one of the most potent human pathogens. According to the latest reports form the World Health Organization (WHO), tuberculosis claimed a mammoth toll of nearly 1.5 million lives globally(1). A hallmark feature attributed to the virulence of this acid-fast bacterium is the unusual composition of its cell wall(2–4). An elaborate, highly hydrophobic capsule and a complex mycomembrane form an additional layer on top of the peptidoglycan and arabinogalactan matrix(5–7). This renders mycobacterial cells impervious to most drug molecules or therapeutic compounds(2, 5, 7, 8). However, in the forefront of determining pathogenic success is a multi-machinery secretion apparatus, called the Type VII secretion system (T7SS), is involved in promoting host cell death, modulation of immune response, zinc and iron assimilation, and maintaining cell wall integrity(5, 9, 10). Essential components of the T7SS include – (i) transmembrane channel and associated membrane proteins that form the cardinal structural element, (ii) AAA+ or FtsK/SpoIIIE ATPases and chaperones that control substrate translocation, (iii) substrates belonging to the PE/PPE family or Esx proteins(9–15). Among the five homologous T7 systems, ESX-1 through ESX-5 (early secretory antigenic target [ESAT-6] secretion system 1-5), ESX-1 is the most well-studied system that is known to secrete at least five virulence associated factors – EsxA (ESAT-6), EsxB (CFP-10), EspB, EspC and EspA(5, 9, 16–18). ESAT-6 and CFP-10 is a WxG100 family of secreted substrate that is encoded by the region of difference (RD1) locus of *M. tuberculosis* genome. It is the absence of this locus that renders compromised virulence in the vaccine strain Bacillus Calmette Geurin (BCG)(9). Similar to the WxG100 family of heterodimeric complexes are the PE/PPE family heterodimeric substrates which have characteristic Pro-Glu(PE) or Pro-Pro-Glu(PPE) motifs in the N-terminal(19). EspB, which is exclusive to the ESX-1 system, is an exception for having the PE and PPE domains joined together by means of a flexible linker in a continuous polypeptide chain(20, 21). It is also the only example of a PE/PPE family secreted substrate that can oligomerise into higher order oligomers, predominantly heptamers(20, 22, 23). Oligomerization of EspB is observed only in the culture filtrate, that is, when the full-length EspB (1-460) is cleaved at the C-terminal domain by the subtilisin-type serine protease MycP_1_(21).

EspB, denoted Rv3881c, is encoded by the extended RD1 region (extRD1) of *Mycobacterium tuberculosis*(24). Several *in vitro* and *in vivo* infection models have previously reported a close association of EspB with host cell toxicity, evasion of phagosome maturation and extrapulmonary dispersal of tuberculosis(24–27). It was shown that the secretion of EspB was unaffected by the co-secretion of other Esp secreted substrates EspA, EspC and EspD(27). Provision of purified C-terminal processed EspB to 5′ Tn:*pe35* mutant, showed elevated levels of toxicity in THP-1 monocytes in a dose dependent fashion(27). This gave rise to a notion that EspB elicits virulence in a manner distinct from EsxA. Crystal structure of monomeric EspB revealed a structured N-terminal domain comprising both WxG and YxxxD signal motif on the same side of a long coiled-coil helix bundle(20, 21). The C-terminal domain, however, remained unstructured due to high structural flexibility. This raises an interesting question – if MycP_1_ cleaves at the C-terminal domain after which the mature isoforms are able to oligomerize – what could be the benefit of retaining ~50 amino acid residue long stretch of disorder? Intriguingly, mature isoform of EspB is able to bind phospholipids such as phosphatidic acid (PA) and phosphatidylserine (PS), while full-length EspB does not show this property(27). Cryo-EM structure of full-length oligomeric EspB illustrated a heptameric ring-like assembly of the N-terminal domain(22). Surprisingly, it was observed that the inner diameter of EspB heptamer was nearly 4.5 nm, which indicated that the central channel was sufficiently large to transport other heterodimeric ESX-1 substrates such as ESAT6-CFP10(22). This notion was further supported when more recently, a 7+1 model of EspB monomer trapped within a heptameric EspB channel was proposed for EspB_2- 348_(23). The structure of EspB_1-460_ and its isoforms has been extensively characterized; however, to date, our understanding of the functional role of EspB remains limited. Therefore, the molecular basis for EspB mediated pathogenesis requires further investigation. Past seminal work by Chen et al, 2013(27), had shown preferential binding of mature EspB with phospholipids nevertheless to date, structural elucidation of this affinity has been elusive. Thus, to provide further insight into the functional aspect of C-terminal processed EspB, we designed a single particle cryo-EM based approach to study EspB-phospholipid interaction in a near-native environment. In this current study, we performed various biochemical and biophysical experiments to characterize the oligomeric states of EspB in the presence and absence of lipid and lipid-membrane. Additionally, we implemented TEM imaging to visualize the EspB interaction with yeast mitochondria and with liposomes, which mimic host outer membrane. Furthermore, interaction of EspB with various lipids were confirmed by MST and leakage assay was performed to show the functional activity of EspB. Our current study provides intriguing evidence of lipid-protein interaction of EspB.

## RESULTS

### Recombinant EspB_1-332_ exists as diverse oligomeric species in solution

Previous reports demonstrated that 1-332 amino acids of EspB of Rv3881c interacts with phospholipids(27). Therefore, the coding sequence corresponding to amino acids 1-332 of Rv3881c was amplified from the genomic DNA of *Mycobacterium tuberculosis* H37Rv and cloned into pET28a vector for heterologous overexpression in *Escherichia coli* (*E. coli*) BL21(DE3) cells (Supplementary Figure 1A). The clone length, as described previously(27) was particularly selected to study the interaction of EspB_1-332_ with phosphatidic acid (PA) and phosphatidylserine (PS) (Supplementary Figure 1A). Despite truncating most of the disordered residues reported for the C-terminal region of the protein, the amino acid sequence indicated a disorder content of nearly 43% while the remaining, predominantly adopted alpha helical structures (Supplementary Figure 1B). Assessment of the alpha fold secondary structure prediction of the construct showed that both termini of EspB_1-332_ contained disordered stretches, approximately ranging between Met1-Asp10 for the N-terminal and residues Ala269-Pro332 for the C-terminal (Supplementary Figure 1C-D). Although high resolution structural characterization could be limited by the presence of such large content of disordered and flexible residues, we proceeded to study EspB_1-332_, resembling secreted EspB, in the presence of phospholipids. At first, recombinant EspB_1-332_ was overexpressed and purified by Ni-NTA affinity chromatography (Figure 1A). The recombinant protein was trypsin digested and was identified to be EspB through MALDI-TOF mass spectrometry (Supplementary Figure 1E). To further improve the homogeneity of purified protein, size exclusion chromatography (SEC) was performed. The gel filtration elution profile was distributed in a majorly trimodal pattern with a small peak separating the two major peaks (Figure 1B). This is in agreement with previous studies citing multiple oligomeric states of EspB(20, 21, 23). For better understanding the oligomeric distribution of EspB_1-332_, the SEC purified fractions were pooled and analyzed with size exclusion chromatography coupled with multi-angle light scattering (SEC-MALS). Interestingly, we observed that the first broad peak obtained in SEC, was split into two peaks, while the other peak profiles remained unaltered in the SEC-MALS elution. We obtained four distinct peaks roughly corresponding to predominant molecular weights of 2.2 MDa, 480 kDa, 251 kDa and 36 kDa, starting from the earliest eluting peak (Figure 1 C). Our results indicate the presence of a broad spectrum of different oligomeric states where the last two peaks hint at the possible heptameric, and monomeric states and the earlier peaks may correspond to higher order oligomeric states. Toward obtaining a visual representation of the nature of oligomers, we took the three peak fractions from SEC, namely Peak 1 at 9 ml, Peak 2 at 11.4 ml and Peak 3 at 13.8 ml to observe the particle distribution under room temperature negative staining transmission electron microscopy (NS-TEM). Our microscopy data revealed intriguing patterns of EspB_1-332_ protein distribution in solution. Peak 1 comprised multiples of ring-like heptamers that associate together in orders of two, three, four and higher to form the first eluting broad fraction observed in SEC (Figure1D, Supplementary Figure 2A). This kind of association of the ring-like EspB_1-332_ also explains the megadalton range of oligomeric mass that we obtained from the SEC-MALS data. Close inspection of the negative staining 2D class averages showed that majorly these rings were attached with their neighbors in a side-to-side fashion (Figure1D, Supplementary Figure 2A). Peak 2 displayed the usual heptameric, homogeneously distributed EspB_1-332_ ring-like oligomers (Figure1D, Supplementary Figure 2B). From the 2D class averages, the ring-like top views and the vase-like side views were distinctly discernible. Therefore, there was no orientation bias observed on the continuous carbon of negative staining TEM grid. Finally, TEM inspection of Peak 3 revealed unique filamentous organizations of EspB_1-332_ approximately ranging between 12-14 nm in length (Figure1D, Supplementary Figure 2C). The open-chain like nature of assembly may explain why this class of oligomers is eluted last in gel filtration, possibly corresponding to monomeric form of EspB_1-332_.

**Figure 1:**
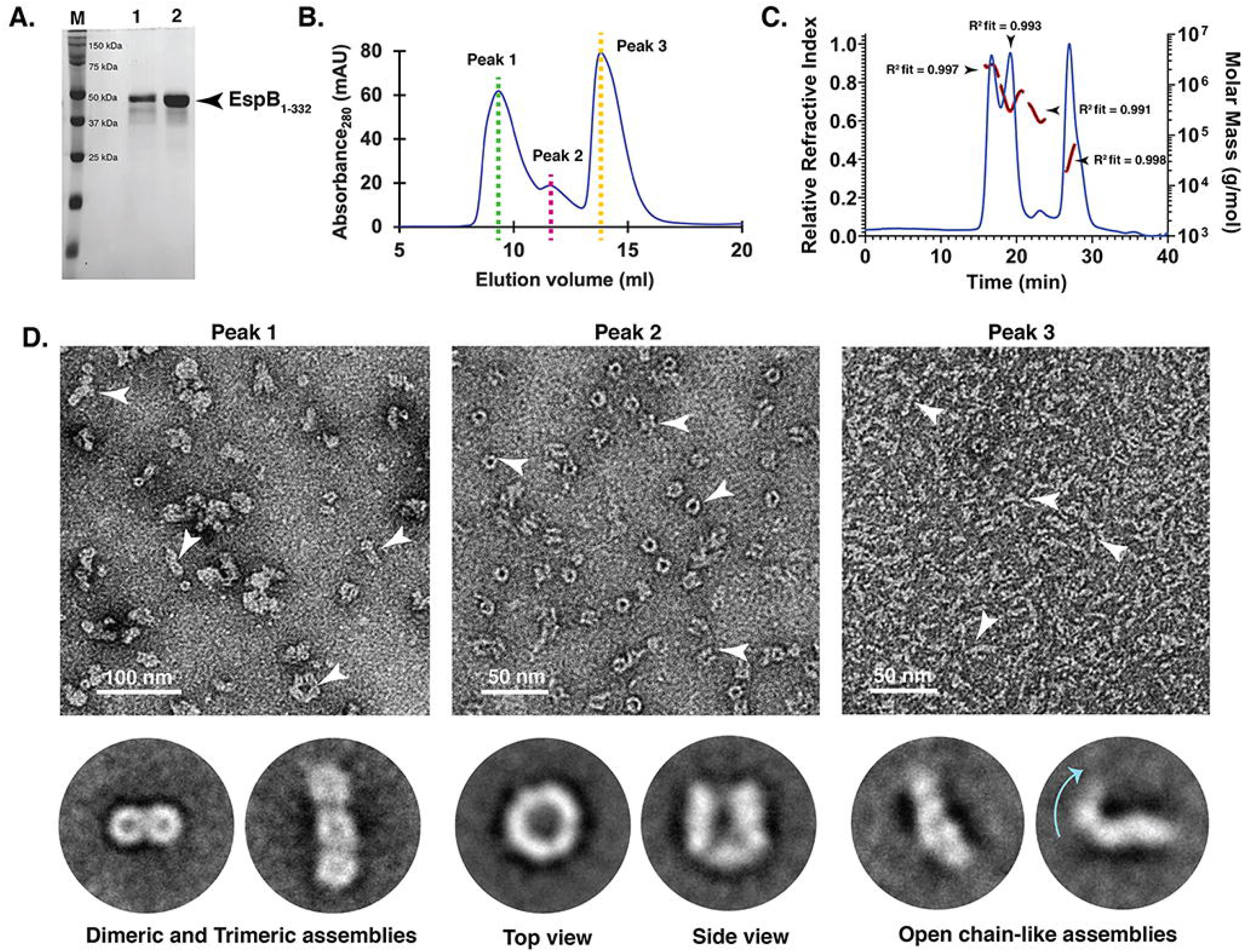
Oligomeric states of purified EspB_1-332_. (A) 12% SDS-PAGE showing the purity of recombinant EspB_1-332_. M – marker, Lane 1 - Ni-NTA purified protein, Lane 2 - SEC purified protein fractions pooled. (B) SEC elution profile shows three broad peaks corresponding to higher order oligomer and plausible monomer population. Green and pink dashed lines show the peak fractions of the high molecular weight assemblies (Peak 1 and Peak 2) of secreted EspB, while yellow dashed line corresponds to the lower molecular weight peak fraction (Peak 3). (C) SEC-MALS profile of EspB_1-332_ obtained after two-step purification. Blue trace denotes refractive index, black trace indicates the molar mass and red trace represents the molar mass fit. (D) Negative staining TEM visualization of the three distinct peak fractions marked in the SEC profile. From left, Peak 1 indicates the presence of multiple fused ring-like oligomers that range from dimers to higher order associations. Bottom panel denotes two representative 2D class averages (enlarged from Supplementary Figure 2A) that show joined rings in the multiples of two and three, respectively. Next, Peak 2 comprises the most homogeneous distribution of oligomeric EspB_1-332_. Representative 2D class averages (enlarged from Supplementary Figure 2B) show the top and side views of ring-like EspB_1-332_. To the right is provided a visualization of Peak 3 that comprises open chain form of EspB_1-332_. Magnified 2D class averages (enlarged from Supplementary Figure 2C) indicate the flexible nature of these chain-like EspB. Cyan arrow marks the area of bending of the protein. White arrowheads have been used to highlight the different oligomeric assemblies in the raw micrographs.

### Single particle cryo-EM 3D reconstruction of purified EspB_1-332_

To evaluate whether the stacking of multiple EspB_1-332_ heptamers could confer upon ESX-1 system its ability to cause virulence in a contact-dependent form, we aimed at performing single particle cryo-EM 3D reconstruction of the conjoined EspB_1-332_ rings. For this, we recorded multiframe cryo-EM movies using a 200 kV Talos Arctica equipped with direct electron detector (DED). However, we consistently observed that the number of joined rings significantly reduced and most of the protein was distributed as free single ring-like particles on the cryo-EM grids (Supplementary Figure 3). These cryo-EM data suggest that EspB multimers (peak 1) were extremely unstable at cryogenic environment and these multimeric interactions were extremely weak, which might easily dissociate at near-physiological cryogenic conditions. Therefore, we did not proceed with extensive data collection because it is possible that even though a significant proportion of EspB_1-332_ exists as fused rings in solution, the stability of these oligomers may be reduced under cryogenic condition and vitreous ice. Next, we sought to obtain structural insights into the discrete heptamers of C-terminal processed EspB_1-332_. Much like full length EspB heptamers, EspB_1-332_ also showed strong preferred orientation on the cryo-EM grids presenting only top views (Figure 2A). This was a stark deviation from the homogeneous orientations observed in our negative staining images (Figure 1D). However, it was interesting to note that the reference-free 2D class averages pointed out a minor subset of hexamers that amounted to nearly 1% of the total particles (Figure 2A). Through cryo-EM we were thus able to identify additional oligomeric stoichiometry among the isolated ring-like EspB_1-332_ population, highlighting the preference of EspB_1-332_ to exist in myriad associations and assemblies. Since orientation bias poses a severe bottleneck in high resolution 3D structure determination, we implemented 0.03% fluorinated octyl maltoside to alleviate particle adsorption at the air-water interface. Consequently, we were able to obtain the multiple orientations of EspB_1-332_ in the presence of surfactant. The uniform distribution of particles across the cryo-EM grid encouraged us to perform 3D reconstruction of EspB_1-332_. After multiple rounds of particle curation, we were able to resolve the structure of EspB_1-332_ at a global resolution of 5.6 Å (Supplementary Figure 4). From our cryo-EM density map, we could distinctly observe the PE and PPE domains harbored in the N-terminal of each EspB_1-332_ monomer. EspB_1-332_ heptamers had an outer diameter of nearly 90 Å and an inner diameter of nearly 45 Å. Thus, truncation of the greater part of the C-terminal domain did not affect the dimensions or N-terminal domain architecture of EspB_1-332_ (Figure 2C). The density for the remaining C-terminal region, which remains unstructured, was undetected in our EspB_1-332_ map thereby reiterating the extent of disorder present. Overall, the current structure shares strong consensus with existing high-resolution characterizations of EspB and therefore it can be considered as a suitable control to study EspB_1-332_ in the context of lipid, which we have discussed in the following sections.

**Figure 2:**
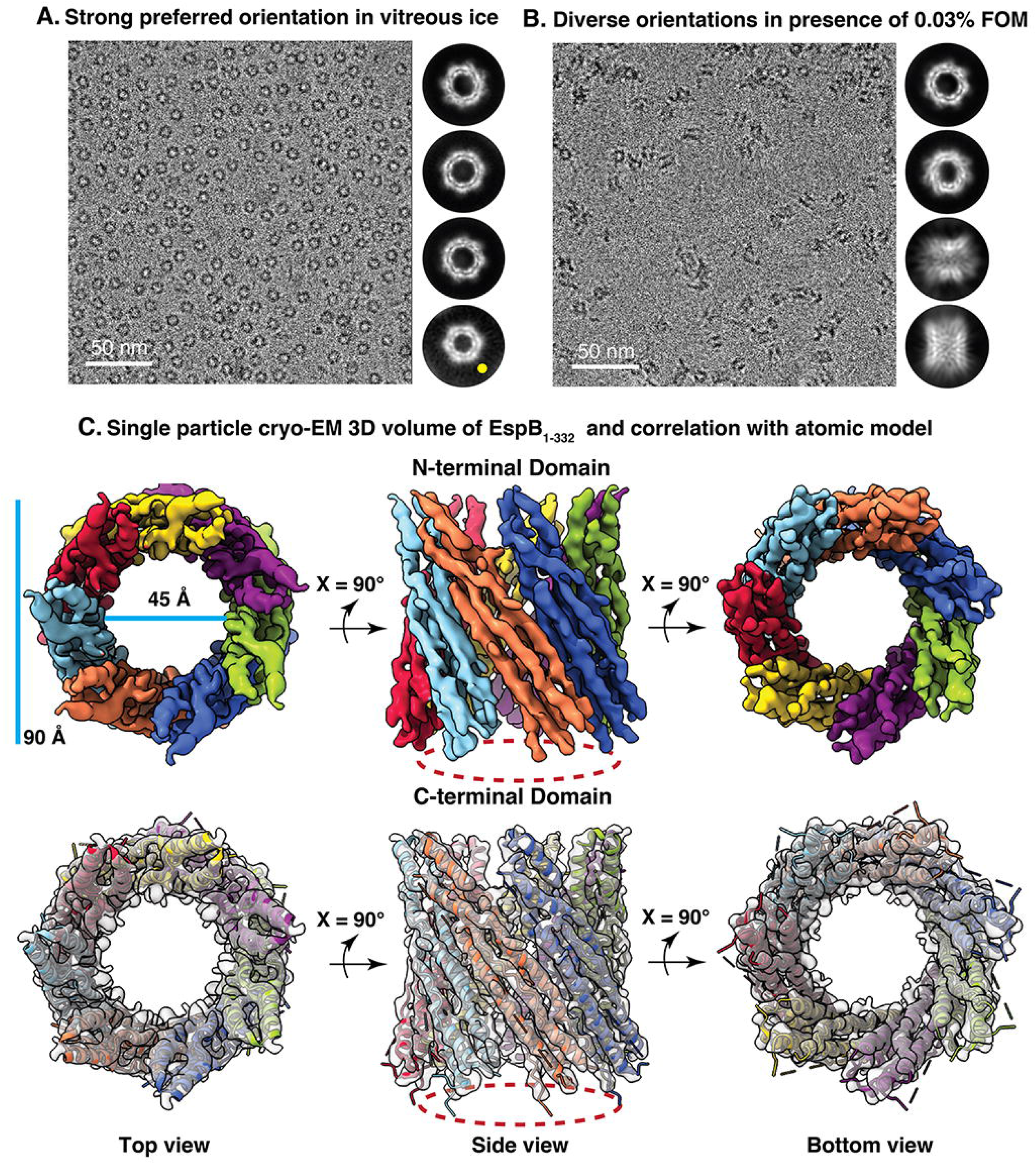
Single-particle cryo-EM structure determination of secreted EspB. (A) Cryo-EM raw micrograph showing a biased orientation distribution of EspB_1-332_ in amorphous ice. Adjacent 2D class averages show only top views of the protein. A minor subset of particles (marked yellow) illustrates the presence of hexameric EspB_1-332_ in a predominantly heptameric population. (B) Cryo-EM raw micrograph represents the effect of ~0.03% fluorinated octyl maltoside on the orientation distribution in amorphous ice. Adjacent 2D class averages show the appearance of side views and tilted along with the previously observed top views. (C) 3D density map of EspB_1-332_ resolved at 5.6 Å reveals strong agreement with the atomic model derived from full-length EspB_1-460_ (PDB ID 6XZC). Upper panel shows the cryo-EM structure where each monomer is coloured differently to highlight the heptameric assembly of the N-terminal domain. The density corresponding to the C-terminal which could not be observed in our map has been traced by dashed red boundary. Transparent rendition of the cryo-EM structure in the bottom panel shows the quality of fitting of the secondary structures into the density map.

### C-terminal processed EspB_1-332_ binds membranes composed of phosphatidic acid (PA)

Chen et al., 2013(27) had proposed that the mature form of EspB, comprising residues 1-332 had a preference to bind lipids like phosphatidic acid (PA) and phosphatidylserine (PS), unlike the full-length EspB_1-460_. However, the way in which secreted EspB can bind PA or PS remains an unanswered question. The presence of a localized stretch of positively charged amino acids in the interior of EspB had invoked a hypothesis that anionic single phospholipids such as PA and PS may be transported through the channel-like N-terminal domain(22). To delve further into the mechanism of phospholipid interaction of C-terminal processed EspB_1-332_, we probed binding with phospholipids - PA and PS, and liposome preparations. At first, we performed liposome sedimentation assay to study membrane interaction of EspB_1-332_ in the context of *E. coli* total cell extract (TCE) liposomes and phosphatidylcholine-cholesterol (PC-Chol) liposomes. Analysis of the resultant supernatant and pellet samples via NS-TEM revealed that the greater proportion of EspB_1-332_ remained in the supernatant while the pellet comprising the liposomes were unassociated with EspB_1-332_ (Figure 3A-B, Supplementary Figure 5). The protein particles that were visible in the pellet fractions were present in the background remaining distinctly excluded from the boundaries of both the TCE liposomes and PC-Chol liposomes. *E. coli* cell membranes are composed of anionic phospholipids, like phosphatidylglycerol (PG) and cardiolipin (CL), therefore the lack of EspB_1-332_ binding to such liposomes concurs with past findings(27). Similarly, PC-Chol liposomes also provide a control to confirm that EspB_1-332_ does not have affinity for PC or Chol. Next, we incubated freshly purified EspB_1-332_ with single phospholipids PA and PS. In the case of PS, we observed that the protein heptamers were homogeneously distributed throughout the micrographs and intermingled with heterogeneously sized clusters of what could be PS molecules (Figure 3C). To confirm the lipid-protein interactions, we prepared two-fold serial dilutions of PS solution and performed microscale thermophoresis (MST) assay with fluorescently tagged EspB_1-332_. Our MST results indicated moderate binding affinity towards PS molecules in the approximate range of K_d_ ~ 6 to 8 μM (Figure 3D). In contrast, for PA treated EspB_1-332_, the NS-TEM micrographs showed unique pattern in which the heptamers were dispersed on the TEM grid. Unlike PS molecules, PA appeared to form small, ordered vesicles and interestingly, the protein particles were prominently localized near or around these lipid structures (Figure 3E). Thus, visual inspection of EspB_1-332_ incubated with PA laid preliminary evidence of a lipid-binding disposition of this esx-1 secreted substrate. This was additionally supported by MST characterization which reported comparatively stronger binding affinity of nearly K_d_ ~ 1 to 4 μM (Figure 3F). Taken together, the tendency of EspB_1-332_ to associate with PA assemblies encouraged us to validate our findings structurally using single-particle cryo-EM.

**Figure 3:**
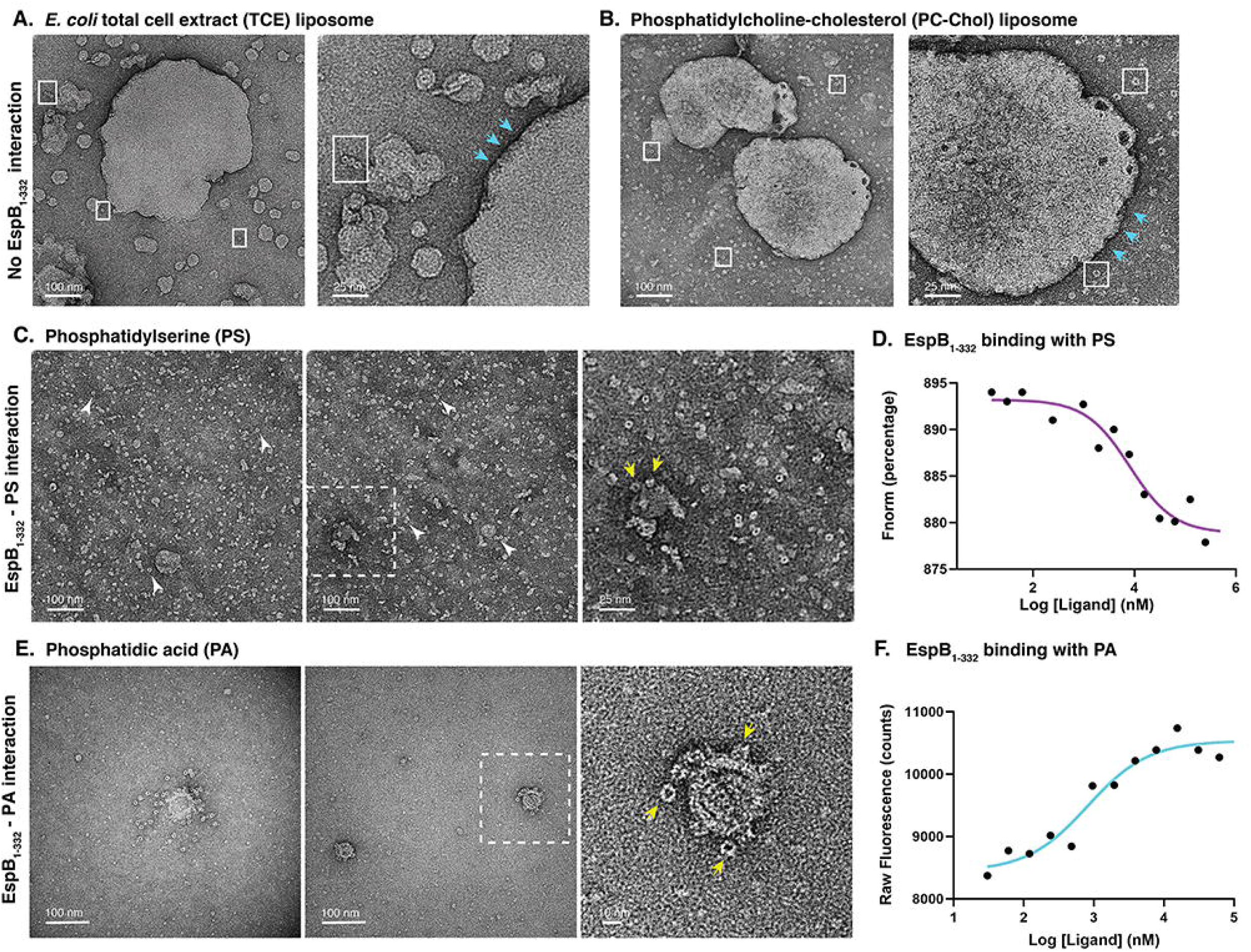
Lipid binding affinity of C-terminal processed EspB_1-332_. (A) A representative negative staining raw micrograph of EspB_1-332_ incubated with liposomes made of *E. coli* total cell extract and enlarged view of the top left corner. White boxes are used to denote the protein and cyan arrows show the boundary of the TCE membrane devoid of any EspB_1-332_ particles. (B) A representative negative staining raw micrograph of EspB_1-332_ incubated with liposomes made of phosphatidylcholine and cholesterol in equimolar ratio and enlarged view of the bottom right corner. White boxes are used to denote the protein and cyan arrows show the boundary of the PC-Chol membrane devoid of any EspB_1-332_ particles. (C) Negative staining raw micrograph of EspB_1-332_ incubated with phospholipid phosphatidylserine. White arrowheads mark free protein and yellow arrows indicate the EspB_1-332_ oligomers that appear to bind plausible lipid architecture. (D) Fluorescence-based binding assay shows micromolar range of binding affinity of EspB_1-332_ with PS molecules, detected in MST mode. (E) Negative staining raw micrograph of EspB_1-332_ incubated with phospholipid phosphatidic acid. Yellow arrows indicate the EspB_1-332_ oligomers that appear to bind plausible lipid architecture. (F) Fluorescence-based binding assay shows micromolar range of binding affinity of EspB_1-332_ with PA molecules, detected in initial fluorescence mode.

**Figure 4:**
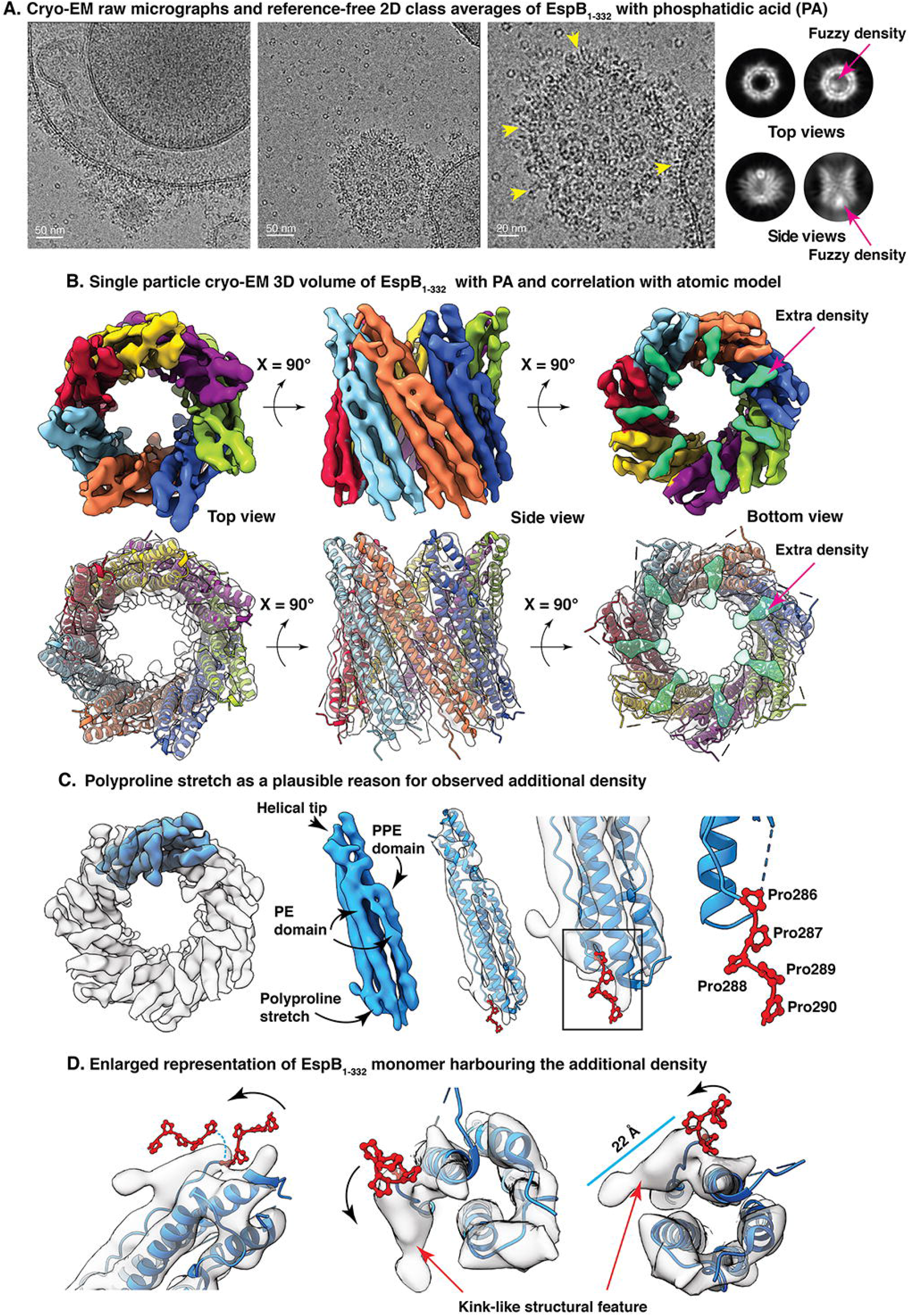
Secreted EspB adheres to membrane bilayer formed by phosphatidic acid. (A) Cryo-EM raw micrographs show a preferential attachment to EspB_1-332_ heptamers on the PA vesicle surface. Yellow arrowheads have been used to pinpoint the side views of the protein attached on the membrane surface. Reference-free 2D class averages show different orientations of the heptamers. Particular top and side views have been highlighted because of the appearance of a fuzzy density at the bottom of the channel. (B) 3D density map of EspB_1-332_ resolved at 6.6 Å indicates coherence with the atomic model derived from full-length EspB_1-460_ (PDB ID 6XZC). Upper panel shows the cryo-EM structure where each monomer is colored differently to highlight the heptameric assembly of the N-terminal domain. The additional density visible in the bottom view of the map has been colored green. Transparent rendition of the cryo-EM structure in the bottom panel shows the absence of modelled coordinates that could fit into the bottom density marked transparent green. (C) Transparent rendition of the electron density map where a single monomer has been colored blue. Monomer derived from the cryo-EM map has been used to show the prominent domains – PE domain, helical tip, PPE domain and the polyproline stretch. Transparent rendition of the monomer fitted with a single EspB chain from PDB 6XZC illustrates the flexible string of multiple prolines. (D) Different orientations of the magnified view of the additional density linked to the helical N-terminal domain. Black curved arrows have been used to show the flexibility of the polyproline stretch. Blue dashed line in the first panel shows a hypothetical trajectory of motion of the prolines to continue into the ~22 Å long extra density which could correspond to the C-terminal domain.

With the objective of exploring secreted form of EspB and PA in a near-native environment, we collected a total of 2601 multiframe cryo-EM movies. The tendency of PA to associate into vesicles remained unaltered under cryogenic conditions and remarkably, we observed the formation of large unilamellar vesicles (LUVs) like structures of various sizes, uniformly occurring throughout vitreous ice (Figure 4A, Supplementary Figure 6). Consistent with our NS-TEM impression, almost all of the EspB_1-332_ heptamers flocked to the PA vesicles leaving very few isolated heptamers in buffer (Figure 4A, Supplementary Figure 6). The vesicles were densely decorated with top views of EspB_1-332_ as well as side views present along the bilayer regions formed by PA. It is also noteworthy to mention that the presence of PA attenuated preferred orientation of EspB_1-332_ alone in amorphous ice. This fortuitous potential of C-terminal processed EspB_1-332_ to be able to specifically adhere to bilayer structures made of PA, urged us to examine the structural changes, if any, upon membrane binding. Toward this goal, we curated an initial set of 751,763 membrane bound EspB_1-332_ particles. Reference-free 2D class averages showed multiple orientations of the protein – top, tilted and side views (Figure 4A). However, we were unable to visualize the distinct bilayer of PA vesicles in the class averages corresponding to the side view of EspB_1-332_ (Figure 4A). Unlike transporters or various pore-forming toxins(28, 29), which embed firmly in between the two leaflets of membrane-bilayer, it is possible that EspB_1-332_, despite having spontaneous binding affinity to PA bilayer, may adhere superficially to the lipid surface. This could potentially lead to averaging out of the signal corresponding to lipid density during the process of particle alignment. Nevertheless, we went ahead with cryo-EM 3D reconstruction of EspB_1- 332_ with PA and were able to obtain the structure at a global resolution of 6.6 Å where the PE and PPE domains in the N-terminal domain were well resolved (Figure 4B, Supplementary Figure 7). Fitting the atomic model of EspB (PDB ID: 6XZC)(22) into our cryo-EM map hinted at strong correlation between the two structures, indicating that PA-binding did not cause any major movement or conformational changes in secondary structures pertaining to the N-terminal region of EspB_1-332_ (Figure 4B). Extending beyond the PPE domain, we observed additional densities in the C-terminal region for which the atomic coordinates have not been modelled (Figure 4B). Further analyzing the connectivity of this ~22 Å long extra density indicated that it may be a continuation of the polyproline stretch that links the N-terminal folded region with the C-terminal unstructured part. (Figure 4C-D). Successively present prolines may potentially induce flexible motion in the loop, allowing it to bend resembling the additional density, thereby leading to the C-terminal domain (Figure 4C-D). However, owing to the high flexibility of disordered residues, application of a uniform B-factor to sharpen the cryo-EM map compromised the connectivity of the dynamic C-terminal region at the given threshold of volume. Particularly in the PA treated dataset, we consistently observed a firm density localized at the bottom of the heptameric ring. This density was present in all the 3D maps including the unfiltered half-maps of the final structure (Supplementary Figure 7). Thus, to improve the interpretability of the structure, we applied a 10 Å gaussian low-pass filter to both the control EspB_1-332_ map as well as to the PA treated structure. On viewing EspB_1-332_ maps at different levels of volume, we observed that the control EspB_1-332_ structure was devoid of any bottom density even at the lowest threshold of density. On the other hand, the appearance of an extra density at the bottom of the PA-EspB_1-332_ map was maintained at all thresholds, getting more pronounced with every step of boosting the volume (Supplementary Figure 8). Consistent with our previous observation, the pronounced bottom density was firmly linked to the N-terminal region (Figure 5A-B). Curiously, this extra density which was exclusive to the PA-EspB_1-332_ map, was located at a position equivalent to where Piton et al. had marked the low-resolution C-terminal domain density in their full-length EspB structure(22) (Figure 5A-B, Supplementary Figure 8).

**Figure 5:**
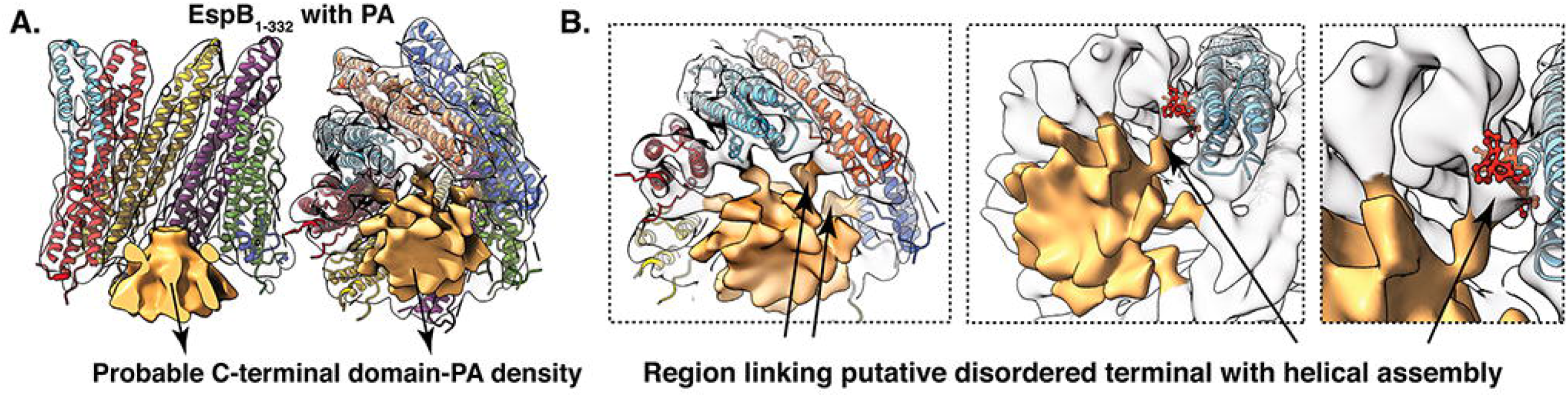
Stabilization of the disordered C-terminal domain in presence of lipid bilayer. (A) Representation of the additional density (marked golden) in the PA-EspB_1-332_ cryo-EM map fitted with PDB 6XZC. (B) Tilted bottom view of the PA-EspB_1-332_ structure shows that the low-resolution bottom extra density is connected to the well-resolved N-terminal domain. Succeeding representations indicate that the proline enriched loop may serve as the link joining the putative C-terminal domain (here, golden density) with the helical N-terminal assembly of EspB_1-332_. Illustrations have been made with a 10 Å Gaussian low-pass filtered version of the PA-EspB_1-332_ map as shown in Figure 4B.

### Characterization of EspB_1-332_ in the context of model membranes comprising PA and PS

Ensuing our observation that EspB_1-332_ possesses membrane bilayer binding property, we sought to study the affinity of EspB_1-332_ towards membranes posing a closer resemblance to host cell environment, in terms of membrane curvature and composition. In most plasma membranes, the outer leaflet mainly consists of PC while our lipids of interest, PA and PS are present in trace amounts. Thus, to provide a more physiologically realizable system, we prepared model liposomes comprising minor proportion of PA and PS, approximately, 54% PC, 29% phosphatidylethanolamine (PE), 5% CL, 5% PA, 5% PS. First, we visualized the model membranes using NS-TEM which showed intact liposomes of diameter of ~100 nm and above (Figure 6A). Having checked the morphology of the control liposomes, we incubated the same with freshly purified EspB_1-332_ heptamers. Surprisingly, we found a biased distribution of the protein particles such that the liposomes were crowded with ring-like EspB_1-332_ molecules with few isolated proteins in the background (Figure 6B). This tendency was maintained all over the NS-TEM grid (Supplementary Figure 9A). We also observed few clusters of EspB_1-332_ around liposomes studded with protein. In some cases, liposomes appeared ruptured with protein top views forming a distinct pattern across the ‘leaky’ areas. Based on TEM imaging alone, it is difficult to comment whether the ruptures were caused by EspB_1-332_ or because of experimental handling. However, it is noteworthy to mention that despite having only 10% of PA and PS combined in the lipid membrane, we were able to obtain very homogenous attachment of EspB_1-332_ over the liposome surface. Thus, even when PC and PE were the main components determining the membrane curvature, the LUV binding ability was not impeded. To obtain a better representation of the protein incubated with our model membrane, we performed reference-free 2D class averaging. As previously obtained, we observed a homogeneous representation of the different orientations of the protein, but few top views and side views showed the presence of extra densities (Figure 6C). Looking through the channel of some top views, we could see fuzzy areas that appear to block the channel from the other end (Figure 6C). Although in negative staining condition we cannot precisely decipher structural information, these classes may hint that the trend of observing additional density at the bottom of EspB_1-332_ when interacting with lipid was conserved. To further confirm the interaction observed through microscopy, we performed binding assay of fluorescently labelled EspB_1-332_ with the liposomes. Overall, the binding affinity was detected in micromolar range with K_d_ ~ 7 to 12 μM, which is more than that recorded for either PA or PS (Figure 6D). Binding assay with other components of the model membrane, PE, PC and CL showed no affinity, thereby indicating that binding of EspB_1-332_ with our model membrane was solely governed by PA and PS (Supplementary Figure 9B-D), even when the vesicles were a mixture of PC, PE, PA, PS and CL. Now, to determine whether the rupturing of liposomes observed in some of the NS-TEM raw micrographs could be attributed to EspB_1-332_, we attempted to estimate membrane permeabilization using carboxyfluorescein release assay. The release of carboxyfluorescein was slow yet consistent reaching ~11.7% in the first 30 mins to ~76.9% with prolonged incubation of 18 hours (Figure 6E). Further, monitoring the percentage leakage over a period of 30 mins, we observed a linear trend (Supplementary Figure 9E). This indicated that EspB_1-332_ may have a role in eliciting virulence by rupturing host cell membrane. It is interesting to note that most pathogens, including *M. tuberculosis* mediate pathogenesis by causing host mitochondrial damage where mitochondrial fission and fusion is severely affected(30–32). Moreover, PA and PS are also known markers of apoptosis. Of particular importance here is PA which regulates intracellular Ca^2+^ concentration(33), one of the primary signals to induce apoptosis(34). Thus, we hypothesized that role of EspB_1-332_ as an esx-1 substrate may be to elicit host mitochondrial damage by binding PA and PS present in the outer cell membrane of mitochondria. To test this hypothesis, we incubated EspB_1-332_ with yeast mitochondria which closely resembles human mitochondria in terms of cell membrane composition. Initially, NS-TEM showed the presence of intact yeast mitochondria where the membrane bilayer appeared continuous (Figure 6F, Supplementary Figure 10A). In the presence of EspB_1-332_, our TEM data revealed distinct localizations of EspB_1-332_ heptamers over the mitochondrial cell membrane (Figure 6G). Regions of broken membrane and clusters of EspB_1-332_ protein were also observed as in the case of incubation with our model membrane (Figure 6B). Notably, this kind of EspB clustering was not visible when only EspB was imaged was NS-TEM. This indicates that exposure to mitochondrial membrane possibly influenced EspB clustering. A particularly interesting observation was the altered morphology of yeast mitochondria post treatment with EspB_1-332_ (Supplementary Figure 10). A significant number of mitochondria appeared to clump together, possibly fusing into large aggregates (Supplementary Figure 10B) while some appeared to be lying on a layer of leaked substances (Supplementary Figure 10C). Thus, through our NS-TEM data we infer that EspB_1-332_ tends to bind yeast mitochondrial membrane and may even impact the intactness of the mitochondrial membrane. However, further cell-based assay is required to validate our TEM and cryo-EM based observations.

**Figure 6:**
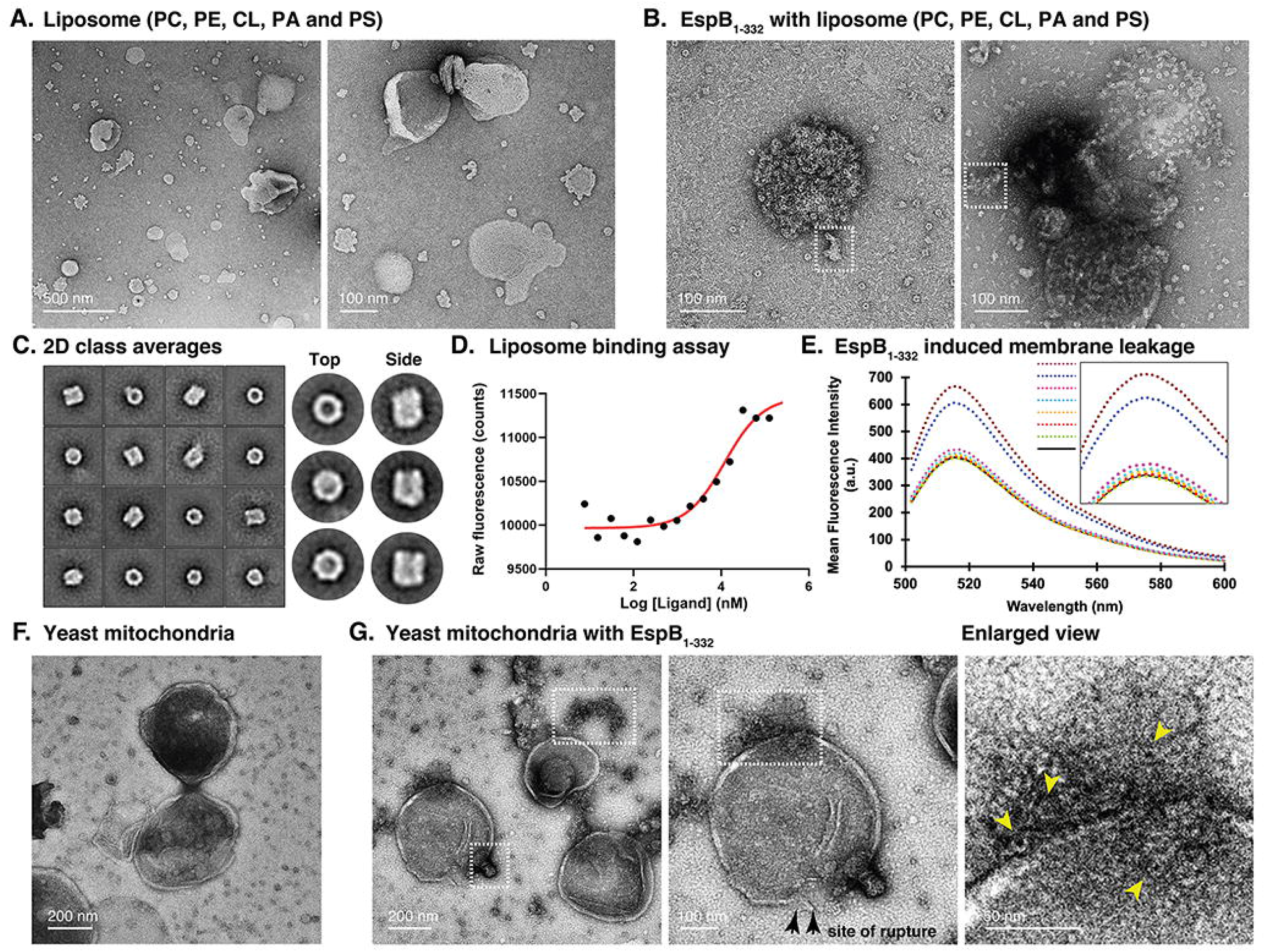
Biophysical characterization of secreted isoform of EspB_1-332_ in the presence of model membranes. (A) Negative staining raw micrographs show liposomes made of PC, PE, PA, PS and CL. (B) Negative staining raw micrograph shows liposomes covered by ring-like EspB_1-332_. White dashed boxes have been used to highlight clustering of EspB_1-332_ around membranes and possible location of bilayer lysis. (C) Reference-free 2D class averages of EspB_1-332_ heptamers incubated with model membrane. (D) Fluorescence characterization of the binding between purified EspB_1-332_ and model liposomes reveals moderate binding affinity. (E) Graph showing over time membrane permeabilization of carboxyfluorescein engulfed liposomes. Black solid curve corresponds to fluorescence of control liposomes. Following dashed curves signify leakage caused by 1.5 μM EspB_1-332_ in time 0 min, 5 mins, 10 mins, 15 mins, 30 mins and 18 hours. Brown dashed curve corresponds to lysis by 0.1% Triton-X. (F) Control *Saccharomyces cerevisiae* mitochondria observed under NS-TEM. (G) NS-TEM raw micrograph showing three isolated mitochondria post incubation with EspB_1-332_. White dashed boxes have been used to highlight clustering of EspB_1-332_ around mitochondrial membrane. A single mitochondrion has been enlarged in the following panels where distinct islands of EspB_1-332_ binding can be observed. Black arrow marks a broken stretch of mitochondrial membrane and yellow arrows highlight ring-like EspB_1-332_ bound at the mitochondrial bilayer surface.

## DISCUSSION

EspB, the largest PE/PPE family substrate secreted by the esx-1 system of *Mycobacterium tuberculosis*, is known to be a key player in mediating host cell pathogenicity. A wealth of structural information shows the architecture of the N-terminal region – PE and PPE domains comprising the YxxxD and WxG motifs respectively, a linker joining the PE/PPE domains and a helical tip(20–23). However, to date, the C-terminal domain remains unstructured. It is thought that the long C-terminal (Pro280-Lys460) causes steric hindrance and prevents the monomers of EspB from oligomerizing within the *M. tuberculosis* cytosol that is why heptameric EspB is undetected in the cell lysate(21). Once EspB_1-460_ is cleaved at Pro332 and A392 by the serine protease MycP_1_(25, 35), mature form of EspB is secreted into the culture filtrate, where it exists predominantly as heptamers. Mutation of Pro332 or Ala392 was shown to be detrimental to the growth of *M. tuberculosis* or *M. marinum*, suggesting an exclusive role of these amino acid residues in Mycobacterial survival(35).Findings by Chen et al, 2013(27), also impressed upon the importance of the C-terminal domain by showing that EspB_1-332_ could selectively bind PA and PS while the EspB_1-460_ with longer stretch of disordered residues did not show lipid binding property. In our current study, we show that EspB_1-332_ binds PA and PS in the context of membrane bilayers. Our cryo-EM structure determination showed that there was an intriguing difference in the C-terminal domain of the EspB_1-332_ control structure and the map obtained when EspB_1-332_ was treated with PA. The control protein sample resembled a hollow ring-like structure comparable to all the previously resolved structures of EspB. However, in the presence of PA, EspB_1-332_ showed a firm density at the location of the disordered C-terminal domain (Figure 5A-B, Supplementary Figure 8)(22), although the C-terminal domain was completely unresolved. The secondary structures that make up the ‘novel’ density in our map are not evident but the overall architecture is reminiscent of the unexplained density that was observed by Solomonson et al. 2015(20) (Figure 5A). Inherent disorder in the protein in association with structural rearrangements of the C-terminal residues may help explain why the resolution of our structure could not be improved further. Additionally, we did not implement mask during refinement of EspB_1-332_ to observe the presence of any extra density associated with PA-EspB_1-332_. This also affects our overall resolution. Despite the possibility that masked-refinement may improve the global resolution, application of a mask around PA-EspB_1-332_ would remove the disordered region and any associated lipid density from the map. This in turn would hamper our target to observe lipid-protein interaction. Close inspection of the PA-EspB_1-332_ structure revealed that this ancillary density was a continuation of the polyproline stretch known to connect the N-terminal helical tip with the low-complexity C-terminal region (Figure 4C-D, Figure 5A-B). Taken together, we infer that this extra density present exclusively in the PA-EspB_1-332_ map could be the disordered C-terminal domain of secreted EspB. This phenomenon of partial structural rearrangements upon membrane interaction appears analogous to how the unstructured N-terminus of alpha synuclein (αSn) adopts a helical secondary structure upon binding negatively charged lipid bilayers(36–38). Similarly, it was shown for ADP-ribosylation factor GTPase activating protein 1 (ArfGAP1) that at low concentration, disorder to order transition of nearly 40 amino acid residues facilitated binding to target membrane(37, 39). Strikingly, the C-terminal of EspB_1-332_ is rich in disorder promoting residues such as alanine, glutamate, glutamine, glycine, lysine, proline and serine(40) (Figure 6B-C). Therefore, it is tempting to consider that interaction with lipid brings order in the otherwise flexible C-terminal domain that is why we were able to observe this region only in the PA dataset and not in the only EspB_1-332_ dataset.

Based on these results, we propose a hypothetical model by which MycP_1_ processed EspB may essay its role as pathogenic substrate of Mycobacterial T7SS system (Figure 7). In solution, EspB oligomers freely tumble and are found in multiple orientations. However, the disordered C-terminal domain cannot be observed in such an environment owing to the dynamic nature of the amino acid residues. Moreover, as the heptameric esx-1 substrate approaches host cell membrane, possibly mitochondrial outer cell membrane, PA and PS affinity ushers EspB to associate with the host membrane. The interaction reduces the degree of freedom of the disordered C-terminal domain and stabilizes the residues thereby making the C-terminal domain visible post membrane binding. Such organization of the C-terminal domain from freely moving to a restricted form may cause mechanical perturbation at the membrane surface causing it to steadily leak the contents from the vesicles. This is supported by our leakage assay that shows relatively slow but steady leakage over time. Thus, we speculate a folding-upon-binding mechanism of EspB mediated cytotoxicity. Further experiments are required to accurately decipher if our hypothesis indeed holds true. However, it is noteworthy to consider that even though MycP_1_ cleaves most of the disordered domain, a sizeable stretch of over 50 disordered amino acid residues is still retained in the secreted isoform of EspB. Although the function of this low-complexity region has not yet been determined, our results combined with past findings indicate that the information to bind membrane is harbored in the C-terminal domain. This may help explain why Korotkova et al.(21), who resolved the crystal structure of EspB_7-278_ in presence of PS, could not obtain the density for PS. Nevertheless, the specificity of C-terminal domain toward lipid binding can be confirmed by truncating the disordered residues, which we intend to investigate extensively in the future. On the other hand, EspB has also been associated with prevention of phagosome maturation(26). Interestingly, phosphoinositides (PIPs) are important in signalling events that lead to phagosome maturation (41). It is possible that EspB_1-332_ may have an affinity for PI (or PIPs), which is another anionic phospholipid like PA and PS, subsequently causing rupture of phagosomal membrane as observed for our model membrane. These ideas pave way for further examination of EspB_1-332_ as a membranolytic esx-1 substrate.

**Figure 7:**
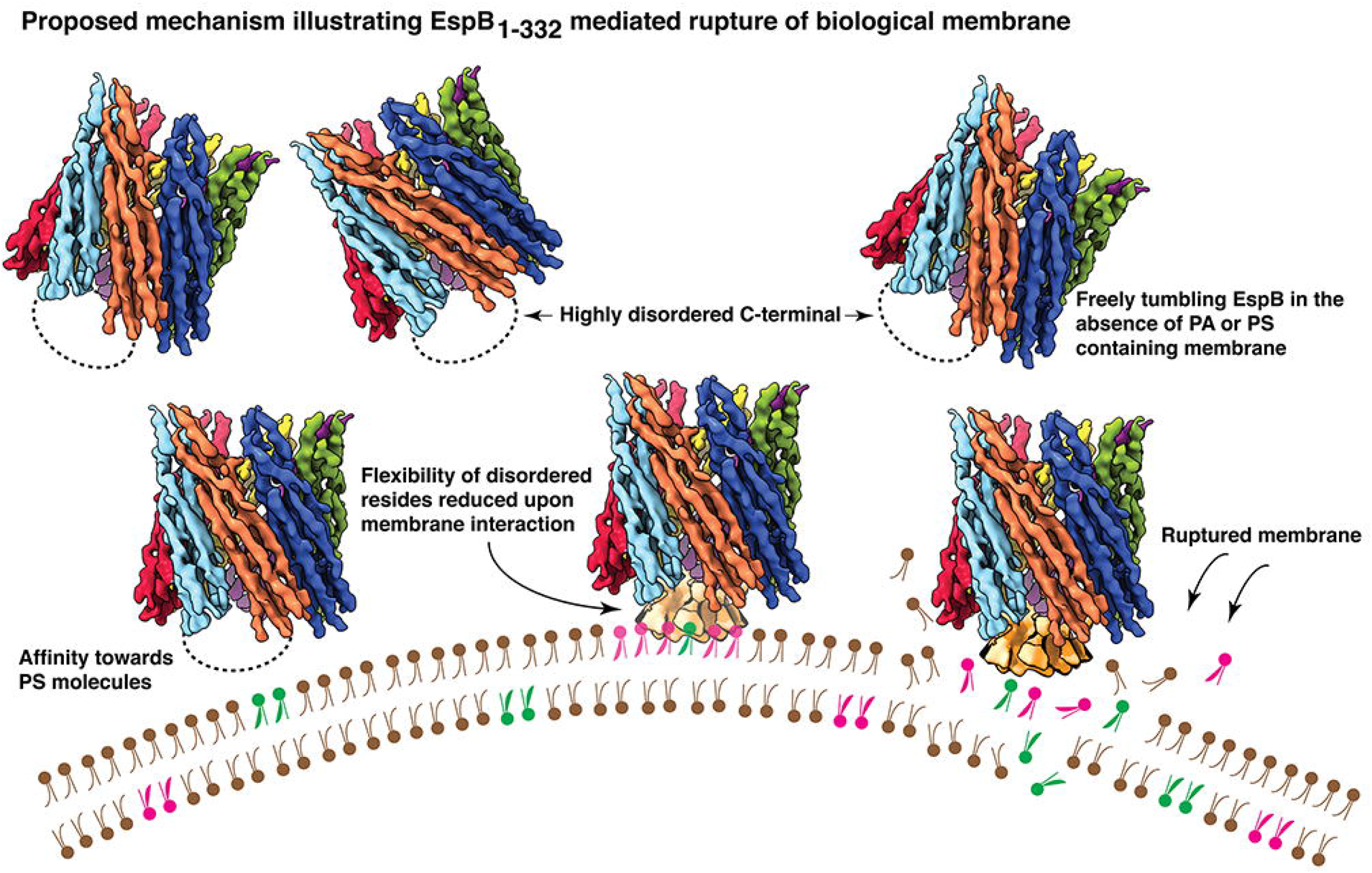
Proposed mechanism of EspB_1-332_ mediated cytotoxicity. Schematic diagram illustrating a hypothetical basis for cytotoxicity caused by MycP_1_ processed EspB. In brief, PA and/or PS binding facilitates disorder to order transition in the low-complexity C-terminal domain. This leads to mechanical perturbance at the membrane surface causing sustained membraned lysis. In figure, pink colour has been used to denote PA molecules and green colour has been used to denote PS molecules.

## Materials and Methods

### Cloning, expression, and purification of EspB_1-332_

The genomic DNA of *Mycobacterium tuberculosis* was a kind gift from Prof. Amit Singh, Indian Institute of Science, Bangalore. Nucleotide sequence corresponding to residues 1-332 of Rv3881c was amplified using Phusion polymerase (NEB) and cloned into pET28a between NdeI and HindIII restriction sites. The clone was confirmed by sequencing and transformed into *E. coli* BL21(DE3) cells for recombinant protein overexpression.

For recombinant protein expression, the *E. coli* BL21(DE3) cells were grown in Luria broth at 37°C to OD_600_ 0.6 and induced for 12 hours with 0.5 mM IPTG at 16°C. Cells were harvested by centrifugation and resuspended in buffer containing 20 mM Tris-HCl pH 8.0, 10 mM Imidazole and 300 mM NaCl. The resuspended cells were lysed by sonication followed by centrifugation at 4°C, 13000 rpm for 1 hour. Recombinant proteins were purified by Ni-NTA affinity chromatography. The cell lysate corresponding to each protein, was loaded onto Ni-NTA column pre-equilibrated with the lysis buffer (20 mM Tris-HCl pH 8.0, 10 mM Imidazole and 300 mM NaCl). The columns were initially washed with the lysis buffer followed by sequential washes containing 20 mM, 40 mM and 80 mM Imidazole. Protein was eluted in a buffer containing 20 mM Tris pH 8.0, 300 mM NaCl and 300 mM Imidazole. During elution, the elute was checked for the presence of protein using Bradford reagent. The protein was analysed for purity in 12% SDS-PAGE gel. This was followed by size exclusion chromatography (SEC) using 24 ml Superdex 200 Increase 10/300 GL column on an AKTA-FPLC system (GE Healthcare). Analytical gel filtration was performed in 20 mM Tris pH 7.5 and 150 mM NaCl at the flow rate of 0.5 ml/min.

### Size exclusion chromatography coupled with multi-angle light scattering (SEC-MALS) of purified EspB_1-332_

The fractions obtained from SEC were pooled and concentrated to ~0.5 mg/ml. SEC–MALS was performed using 20 mM Tris pH 7.5 and 150 mM NaCl buffer equilibrated analytical Superdex 200 Increase 10/300 GL gel filtration column (GE Healthcare) on a Shimadzu HPLC. Protein peaks resolved after size exclusion, were subjected to in-line refractive index (Waters Corp.) and MALS (mini DAWN TREOS, Wyatt Technology Corp.) detection to estimate the molar mass. The data acquired from UV, MALS and refractive index (RI) were analyzed using ASTRA 6.1 software (Wyatt Technology).

### Preparation of lipids and liposome sample

#### PA and PS preparation for microscopy and binding study

Powder form of 3-sn-phosphatidic acid (Sigma Aldrich), Brain phosphatidylserine (Avanti Polar Lipids) was dissolved in chloroform in a clean glass vial and evaporated in a desiccator overnight at room temperature. Thin film of dried lipid obtained after drying was dissolved in 20 mM Tris pH 7.5 and 150 mM NaCl and stored in −20°C for future use.

#### Liposome preparation

##### TCE liposome

Powder form of *E. coli* Total Cell Extract (Avanti Polar Lipids) was dissolved in 20 mM Tris pH 7.5 and 150 mM NaCl and incubated at 55°C for 30 mins. This colloidal heated solution was subjected to extrusion (Mini Extruder, Avanti Polar Lipids) using 200 nm pore-sized polycarbonate membranes. Using the Avanti polar extruder apparatus, the colloidal solution was passed through the polycarbonate membranes for a total of eight rounds at 55°C. The resultant clearer solution containing unilamellar vesicles were used for downstream experiments and the remaining was stored at 4°C for not more than 3 days.

##### PC-Chol liposome

Powder form of Egg-PC (Sigma Aldrich) and cholesterol (Sigma Aldrich) was dissolved in dissolved in chloroform. Next, 250 μM chloroform solubilised PC was mixed with 250 μM chloroform solubilised cholesterol and kept for overnight evaporation. The thin lipid film of mixed lipids obtained after removal of residual chloroform was dissolved in 500 μl of 20 mM Tris pH 7.5 and 150 mM NaCl with gentle pipetting. The solution was next incubated at 55°C for 30 mins. This colloidal heated solution was subjected to extrusion as described above. The PC-Chol unilamellar vesicles were used for downstream experiments and the remaining was stored at 4°C for not more than 3 days.

##### Model membrane preparation

Powder form of Egg-PC (Sigma Aldrich), L-α-phosphatidylethanolamine (Sigma Aldrich), 16:0 Cardiolipin (Avanti Polar Lipids), 3-sn-phosphatidic acid (Sigma Aldrich), Brain phosphatidylserine (Avanti Polar Lipids) was dissolved in chloroform. Next, 270 μM PC, 145 μM PE, 25 μM CL, 25 μM PA, 25 μM PS were mixed and kept for overnight evaporation. The thin lipid film obtained the following day, after removal of residual chloroform was dissolved in 500 μl of 20 mM Tris pH 7.5 and 150 mM NaCl with gentle pipetting. The solution was next incubated at 55°C for 30 mins. The resultant colloidal heated solution was subjected to extrusion as mentioned above. Finally, the model membrane liposomes comprising a mixture of PC, PE, CL, PA and PS were were used for downstream experiments and the remaining was stored at 4°C for not more than 3 days.

##### Carboxyfluorescein (CF) engulfed liposomes

Liposomes comprising CF were prepared following previously described methodology(42). Briefly, at first a 20 mM CF stock solution (pH 7.4) was prepared. For largescale preparation of liposomes, chloroform dissolved solution of 2mM PC, 1.2 mM PE, 0.2 mM CL, 0.2 mM PA and 0.2 mM PS were mixed and kept for overnight evaporation. The the resultant dried, thin lipid film was dissolved with gentle pipetting in 500 μl of 20 mM Tris pH 7.5 and 150 mM NaCl. To this solution, 500 μl of 20 mM CF solution was added and mixed by gentle pipetting. This was followed by incubation at 55°C for 30 mins. Next, the mixture was sonicated in a bath sonicator three times for 30 secs with intermittent vortexing for 15 secs. This was followed by extrusion according to steps mentioned above. Following extrusion, the liposomes that harboured CF were separated from free-CF by using a room-temperature gravity sizing column pre-packed with Sephadex G-25 resin. The CF-liposomes were then stored at 4°C and used the following day for membrane permeabilization assay.

#### Microscale thermophoresis (MST)

EspB_1-332_ and free phospholipid or model membrane interaction assay was done using MST (Nanotemper)(43). Freshly purified EspB_1-332_ was labelled with RED-NHS 2^nd^ generation amine reactive dye equimolar ratio. Monolith™ NT.115 standard MST capillaries were used in each experiment. In binding reaction, the labelled protein concentration was kept constant at approximately 20 nM. The protein was incubated with 16 two-fold serial dilutions of ligands dissolved in 20 mM Tris (pH 7.5), 150 mM NaCl. For PA, PS, CL, PC and PE, the highest ligand concentration was 500 μM and the highest concentration for the model membrane was 250 μM. The protein and lipids/membrane were both diluted in 20 mM Tris (pH 7.5), 150 mM NaCl and 0.05% Tween-20. The assays were performed with two independent set of purifications and each time binding with PA, PS and model membrane was obtained in micromolar range. The samples were analyzed with a Monolith™ NT.115 pico device using MO.Control Software (Nanotemper). For PA and model membrane, binding was detected in the initial fluorescence mode and for PS the binding was recorded in the MST mode on time 10 secs.

#### Liposome leakage assay

The fluorescence due to only CF-liposomes were recorded at an emission wavelength of 512 nm upon excitation at 492 nm, using Cary Eclipse spectrophotometer. The CF-liposomes were incubated at room temperature with 1.5 μM freshly purified EspB_1-332_ protein and the fluorescence was recorded at intervals – 0 min, 5 mins, 10 mins, 15 mins, 30 mins. The reaction mixture was further incubated at room temperature and the next reading was taken post 18-hour incubation. As control to signify 100% lysis, 0.1% Triton-X was added to the CF-liposomes. Experiment was performed in triplicates. Percentage leakage at a specific time was calculated using the following formula:

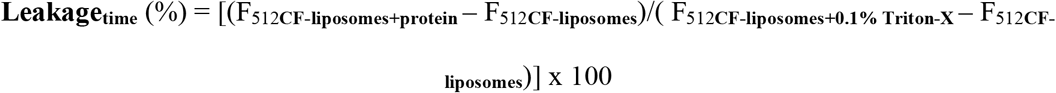

#### Negative staining transmission electron microscopy (NS-TEM) sample preparation

##### TEM analysis of purified EspB_1-332_

For each peak fraction (9 ml, 11.4 ml and 13.8 ml), approximately 3.5 μl of 0.1 mg/ml protein was added on 400-mesh Cu TEM grids which were freshly glow-discharged (negative polarity) for 30 secs in GloQube glow-discharge system. The sample was incubated at room temperature for 1 minute. The surplus solution was then carefully blotted off using Whatman filter paper. This was followed by negative staining using 1% freshly prepared uranyl-acetate solution.

##### TEM analysis of heptameric EspB_1-332_ with TCE liposomes, PC-Chol liposomes

Approximately 1 mg/ml EspB_1-332_ heptamers were incubated with TCE and PC-Chol liposomes at room temperature for 20 mins. The reaction mixtures were then subjected to centrifugation at 13,000 rpm for 30 mins (liposome sedimentation assay). The supernatant was separated from the pellet. The pellet comprising the bulk of the liposomes were dissolved in 20 mM Tris (pH 7.5), 150 mM NaCl buffer such that the volume was equal to that of the supernatant. The supernatants and the pellet solutions were negatively stained as described above.

##### TEM analysis of heptameric EspB_1-332_ with PA and PS

Approximately 1 mg/ml EspB_1-332_ heptamers were incubated with 1 mM PA and PS solutions respectively, and incubated at room temperature for 20 mins. This was followed by negative staining as described above.

##### TEM analysis of heptameric EspB_1-332_ with model membrane

Approximately 1 mg/ml EspB_1-332_ heptamers were incubated with model membrane solution and incubated at room temperature for 20 mins. This was followed by negative staining as described above.

##### TEM analysis of heptameric EspB_1-332_ with yeast mitochondria

*Saccharomyces cerevisiae* mitochondria was a kind gift from the laboratory of Prof. Patrick D’Silva, Indian Institute of Science, Bangalore. Approximately 0.5 mg/ml EspB_1-332_ heptamers were incubated with 50 ng/μl yeast mitochondria at room temperature for 20 mins. This was followed by negative staining as described above.

The TEM data were acquired on a 120 kV Talos L120C room temperature electron microscope equipped with a bottom-mounted Ceta camera (4Kx4K) at different magnifications ranging between 43,000x – 120,000x magnification between calibrated pixel size of 3.68-0.92 Å/pixel at specimen level.

#### Negative staining transmission electron microscopy (NS-TEM) data processing

The negative staining TEM raw micrographs were imported into EMAN 2.1(44) and the best micrographs were retained for calculating the reference-free 2D class averages. Around 3000 particles were manually picked for EspB_1-332_ fused rings, EspB_1-332_ heptamers, EspB_1-332_ monomers and EspB_1-332_ heptamers with model membrane using EMAN 2.1, and particles were extracted using e2boxer.py in EMAN 2.1. The extracted particles were imported into RELION 3.0(45) and two to three rounds of reference-free 2D class averages were calculated to classify the particles to increase the signal to noise ratio. The class averages with prominent features were selected for EspB_1-332_ fused rings, EspB_1-332_ heptamers, EspB_1-332_ monomers and EspB_1-332_ heptamers with model membrane samples respectively. The cleaned particle sets thus obtained after the initial rounds of 2D classification, were re-classified using simple_prime2D of SIMPLE 2.0(46).

#### Cryo-electron microscopy (cryo-EM) sample preparation and data collection

The EspB_1-332_ heptamer protein sample was taken for cryo-electron microscopy to study with and without PA. Initially, 300 mesh copper Quantifoil R 1.2/1.3 grids were glow discharged in GloQube glow discharge apparatus at 20 mA for 90 seconds. For control EspB_1-332_ heptamers, protein was concentrated to 6 mg/ml and 0.03% fluorinated octylmaltoside (FOM) was added to the sample immediately before application on cryo-TEM grid. 3 μl of the FOM-EspB_1-332_ sample was applied to the glow discharged grids and incubated at 100% humidity for 10 secs. Excess sample was blotted for 7 seconds at −2 blot force.

In the case of EspB_1-332_ heptamers in presence of PA, nearly 1 mg/ml protein was incubated with 2 mM PA at room temperature for 20 mins. 3 μl of the PA-EspB_1-332_ sample was applied on the glow discharged grids and incubated at 100% humidity for 10 secs. Excess sample was blotted for 7 seconds at 0 blot force.

This was followed by plunge-freezing in liquid ethane cooled by ambient liquid nitrogen in FEI Vitrobot Mark IV plunger. Imaging was performed in a Thermo Scientific™ 200 kV Talos Arctica Transmission Electron Microscope equipped with K2 Summit Direct Electron Detector (4Kx4K) (Gatan Inc). Data collection was performed using automated data collecting software package, LatitudeS(47) (Gatan Inc.) at 54,000x magnification at a calibrated pixel size of 0.92 Å/pixel at specimen level. Movies were recorded for 40 frames and 8 secs with a total calibrated dose of 60 e^−^/Å^2^. The defocus range for data collection ranged from −0.75 μm to −2.5 μm.

#### Cryo-electron microscopy (cryo-EM) data processing

For both the cryo-EM datasets, the same steps were followed as mentioned next. A total of 3000 movies were first corrected for beam-induced motion using RELION 3.1 implementation of MotionCor2(48). The resultant motion corrected micrographs were imported into cisTEM(49) to sort the data based on the signal to noise ratio and to select micrographs with signal better than 7 Å resolution. The contrast transfer function (CTF) of the selected micrographs was estimated using CTFFIND 4.1.13(50). A small subset of particles were manually picked in RELION 3.1 to generate the reference-free 2D class averages. The best 2D classes were selected to use as a template for automated particle picking.

##### Control EspB_1-332_ data processing pipeline

Approximately 630375 particles were picked and extracted at a box size of 240 pixels. After two rounds of reference-free 2D classification, 589429 particles were obtained. These curated particles were next subjected to a round of 3D classification using C7 symmetry. The best class showing the highest resolution and distinct structural features was selected for autorefinement with C7 symmetry. The contrast transfer function of the refined particle set was further improved using beamtilt estimation followed by a round of autorefinement using C7 symmetry. The final cryo-EM density map of control EspB_1-332_ heptamer was obtained at 5.6 Å at 0.143 Fourier Shell Correlation (FSC).

##### PA-EspB_1-332_ data processing pipeline

Approximately 751763 particles were picked and extracted at a box size of 240 pixels. After two rounds of reference-free 2D classification, 300140 particles were obtained. These curated particles were next subjected to a round of 3D classification using C7 symmetry. The class with high resolution structural feature was selected for autorefinement using C7 symmetry. Beamtilt estimation of the refined particle set did not further improve the map resolution. The final cryo-EM density map of PA-EspB_1- 332_ heptamer was obtained at 6.6 Å at 0.143 Fourier Shell Correlation (FSC).

All visualizations were performed using UCSF Chimera(51) and UCSF ChimeraX(52).

## ACKNOWLEDGEMENTS

We acknowledge Department of Biotechnology, Department of Science and Technology (DST) and Science, and Ministry of Human Resource Development (MHRD), India for funding and cryo-EM facility at IISc-Bangalore. We acknowledge DBT-BUILDER Program (BT/INF/22/SP22844/2017) and DST-FIST (SR/FST/LSII-039/2015) for National Cryo-EM facility at IISc, Bangalore. We acknowledge the financial support from the Ministry of Human Resource Development (MHRD) (Grant Number-STARS-1/171), and DBT (Grant No. BT/PR25580/BRB/10/1619/2017) for financial support. We thank DBT-IISc partnership program for TEM facility at Biological Sciences Division. We would like to acknowledge the LC-MS facility and SPR facility at Biological Sciences Division for acquisition of the mass spectrometry analysis and binding affinity characterization studies. We are grateful to Dr. Gopinath Chattopadhyay for his help with the binding studies.

## AUTHOR CREDIT STATEMENT

Nayanika Sengupta – Conceptualization, Investigation, Formal Analysis, Methodology, Visualization, Writing – original draft and editing.

Surekha P – Investigation (cryo-EM raw micrographs screening and particle curation for both cryo-EM datasets), Writing – reviewing and editing.

Somnath Dutta – Conceptualization, Supervision, Formal Analysis, Resources, Visualization, Writing – original draft, reviewing, editing and funding.

## DATA AVAILABILITY

The cryo-EM density maps corresponding to EspB_1-332_ and EspB_1-332_ in the presence of PA, have been deposited to Electron Microscopy Data Bank (EMDB).

