## Supplemental Figure for "Cryo-EM reveals the membrane binding phenomenon of EspB, a virulence factor of the Mycobacterial Type VII secretion system"

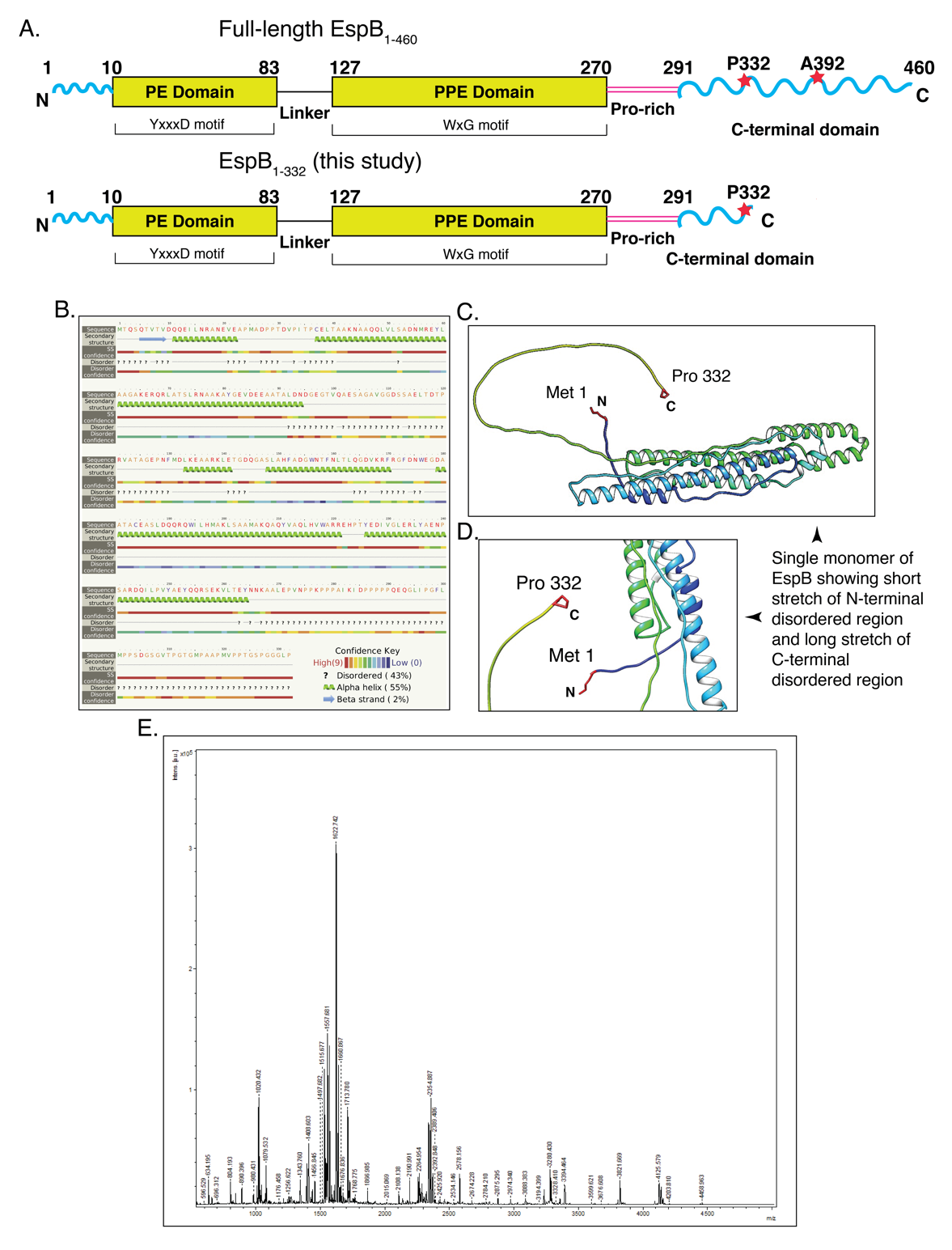

**Supplementary Figure 1**: (A) Schematic representation of the different domains harbored in full-length EspB_1-460_ and in EspB_1-332_ used in this study. Blue curved lines represent areas of disorder, rectangular boxes represent the folded PE and PPE domains, magenta-colored parallel lines show the proline rich region that links the N-terminal domain to the C-terminal domain. Red stars denote MycP_1_ cleavage sites. (B) Secondary structure prediction of EspB_1-332_, using Phyre2 showing a majorly α-helical N-terminal domain and a disordered C-terminal domain. (C) Alpha fold representation of monomeric EspB_1-332_ denoting the relative orientation of the N-terminal Met and C-terminal Pro. (D) Enlarged view of the flexible terminal of EspB_1-332_ reveals a short N-terminal disorder stretch and a nearly ~50 residue long disordered C-terminal. (E) MALDI-TOF spectrum of recombinant EspB_1-332_.

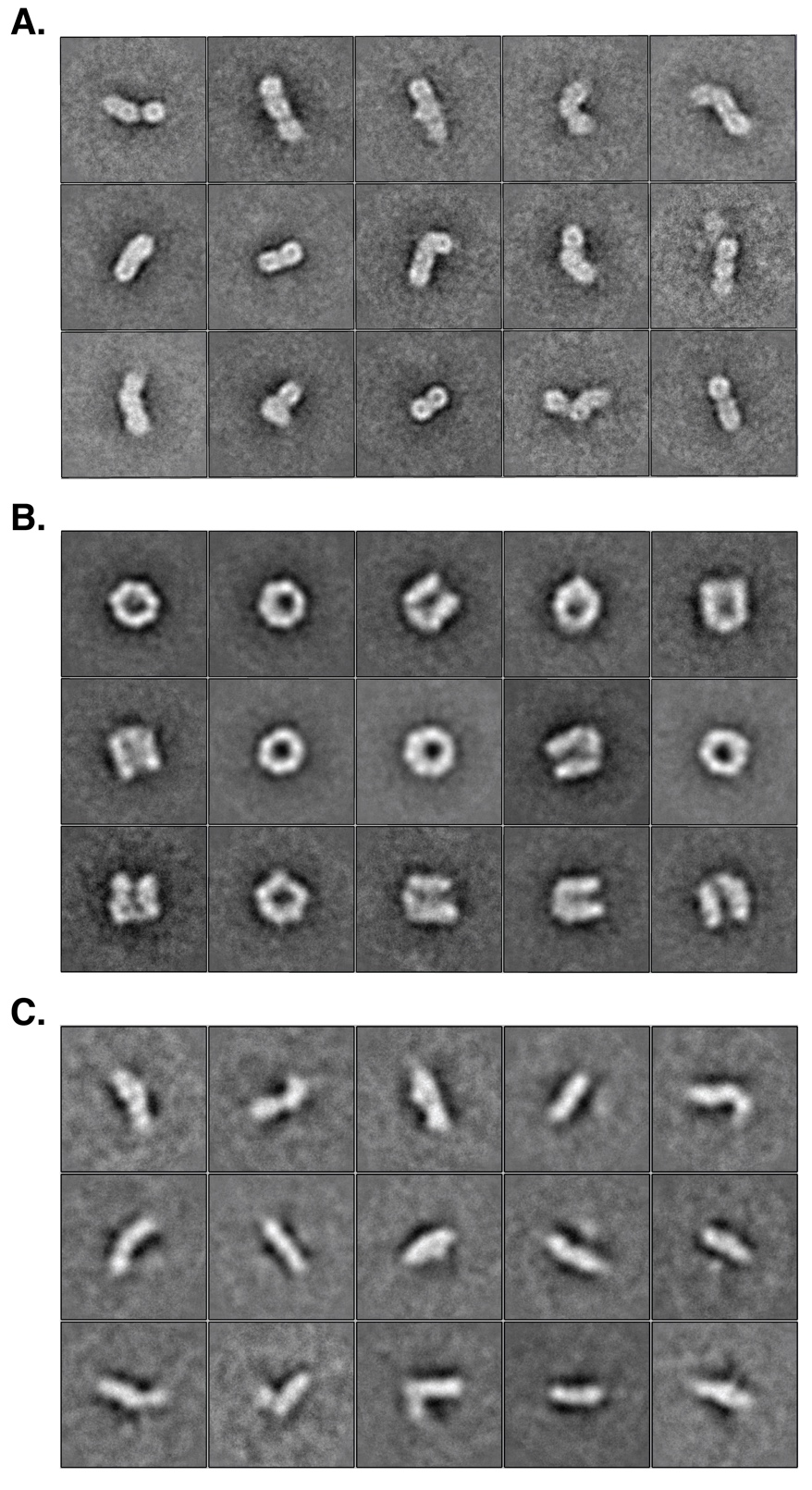

**Supplementary Figure 2**: **Negative staining transmission electron microscopy (NS-TEM) 2D class averages.** (A) Extended 2D classes of the higher order oligomeric fraction of EspB_1-332_, selected views of which have been listed in Figure 1D. (B) Extended 2D classes of the discrete ring-like population of EspB_1-332_, selected views of which have been listed in Figure 1D. (C) Extended 2D classes of the open chain-like population of EspB_1-332_, selected views of which have been listed in Figure 1D.

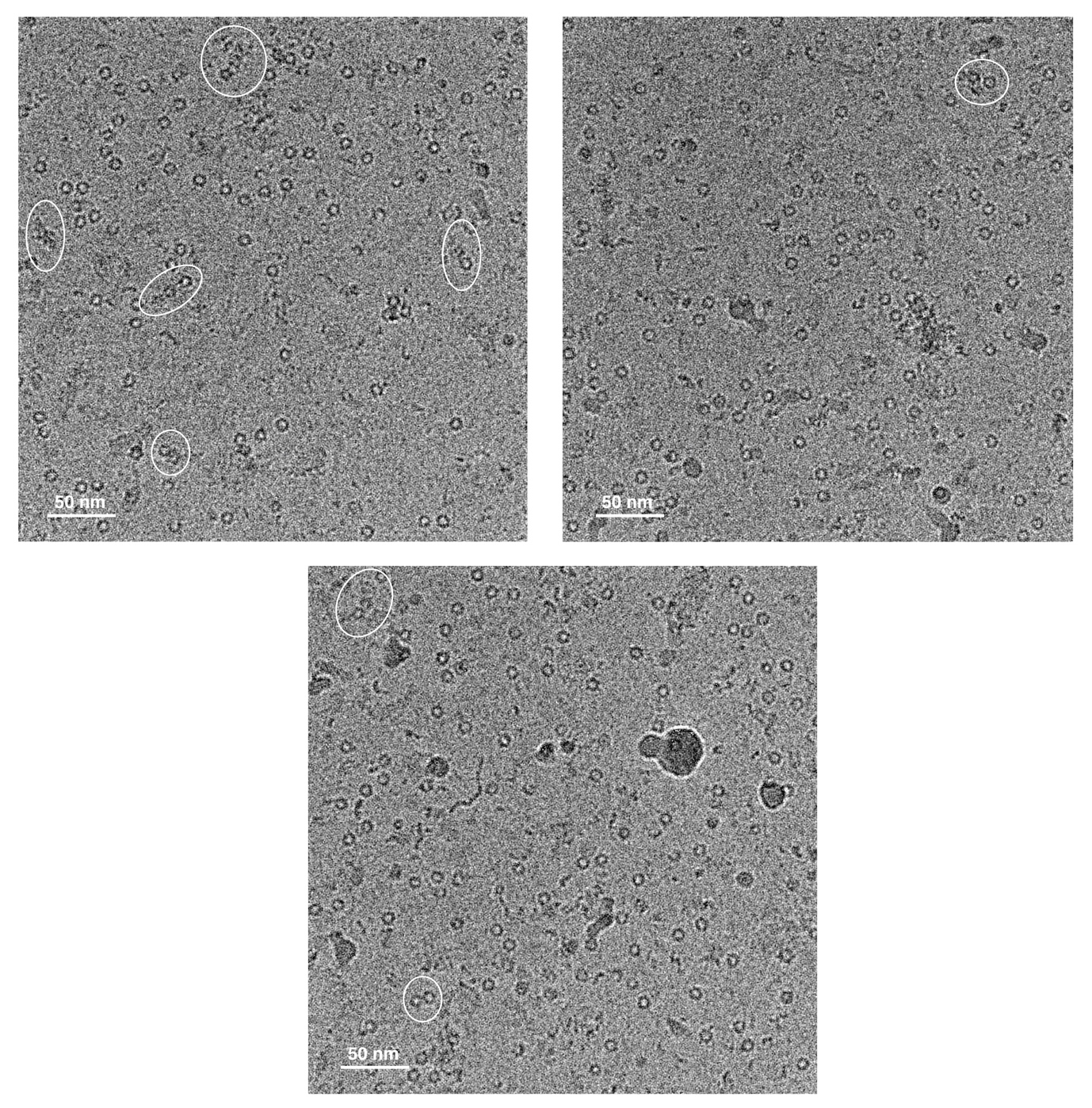

**Supplementary Figure 3**: **Different areas of the cryo-EM grid reveals the instability of the fused EspB_1-332_ rings in cryogenic conditions.** Major population of the multimers as observed in NS-TEM (Figure 1D) appear to be separated into individual hexameric or heptameric rings. Few conjoined oligomers coexist with ring-like oligomer and have been demarcated within white boundaries.

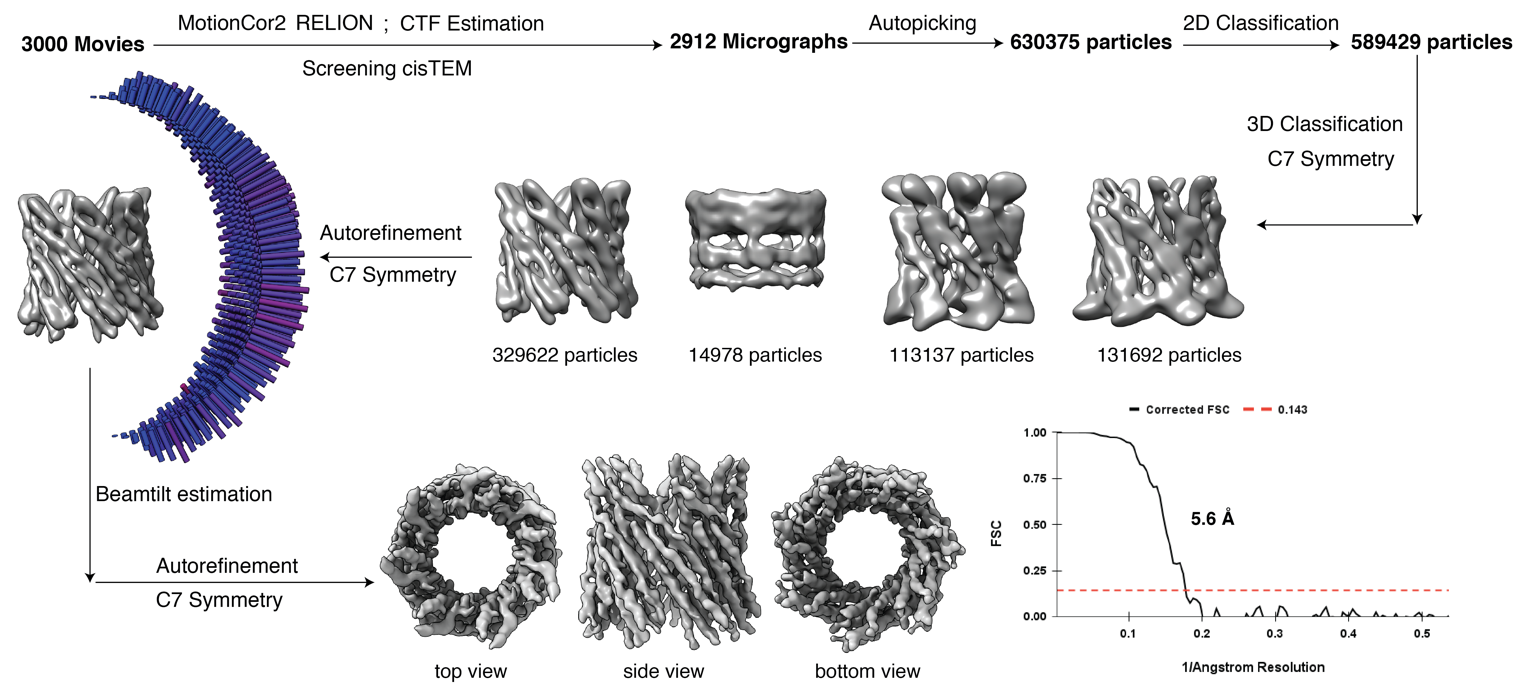

**Supplementary Figure 4**: Cryo-EM data processing pipeline followed for control EspB_1-332_ dataset. Gold standard Fourier Shell Correlation (FSC) calculation shows a resolution of 5.6 Å. See Methods for extended procedures.

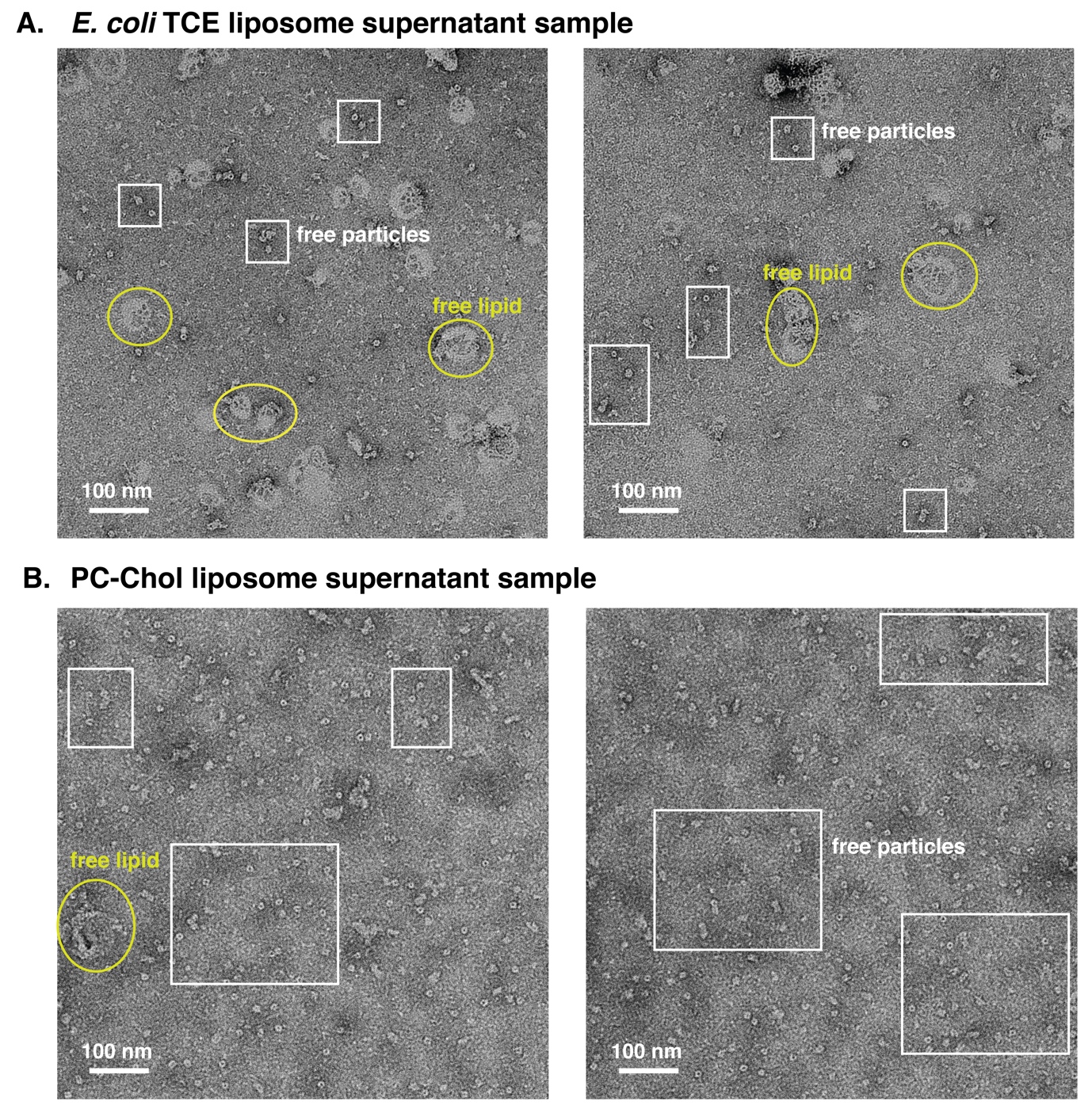

**Supplementary Figure 5**: **NS-TEM analysis of the supernatant fractions obtained after liposome sedimentation assay.** (A) *E. coli* TCE liposome treated EspB_1-332_ predominantly segregates into the supernatant as compared to the pellet (Figure 3A). Proteins appear isolated from the light stained areas which possibly denote lipid patches. (B) PC-Chol liposome treated EspB_1-332_ predominantly segregates into the supernatant as compared to the pellet (Figure 3B). White boxed areas show free particles of protein whereas the yellow elliptical boundaries demarcate the free lipids.

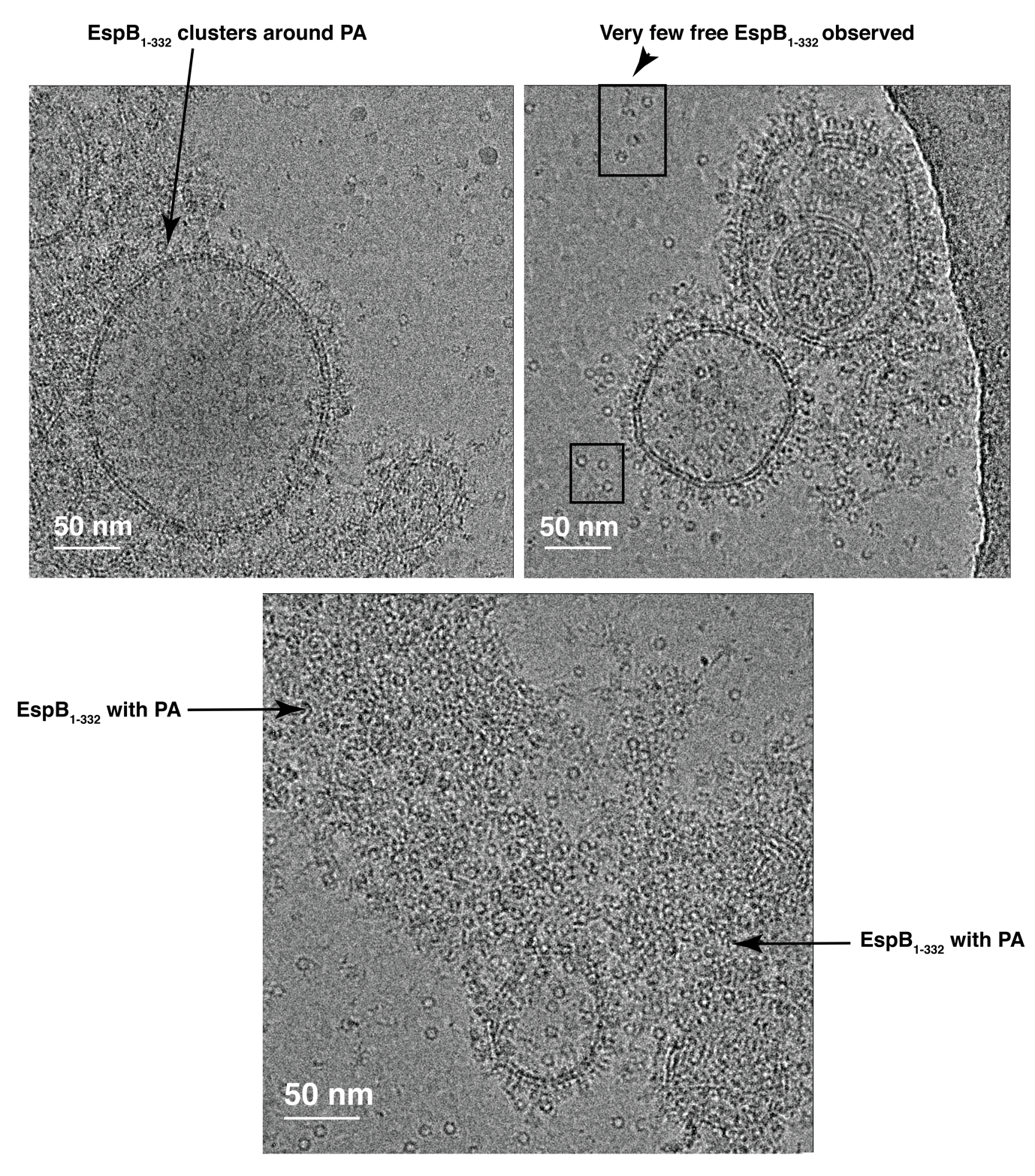

**Supplementary Figure 6**: **A montage of cryo-EM raw micrographs that show a remarkable affinity of EspB_1-332_ towards PA vesicles.** Only few protein particles appear in the background while most of the EspB_1-332_ cluster around PA vesicles.

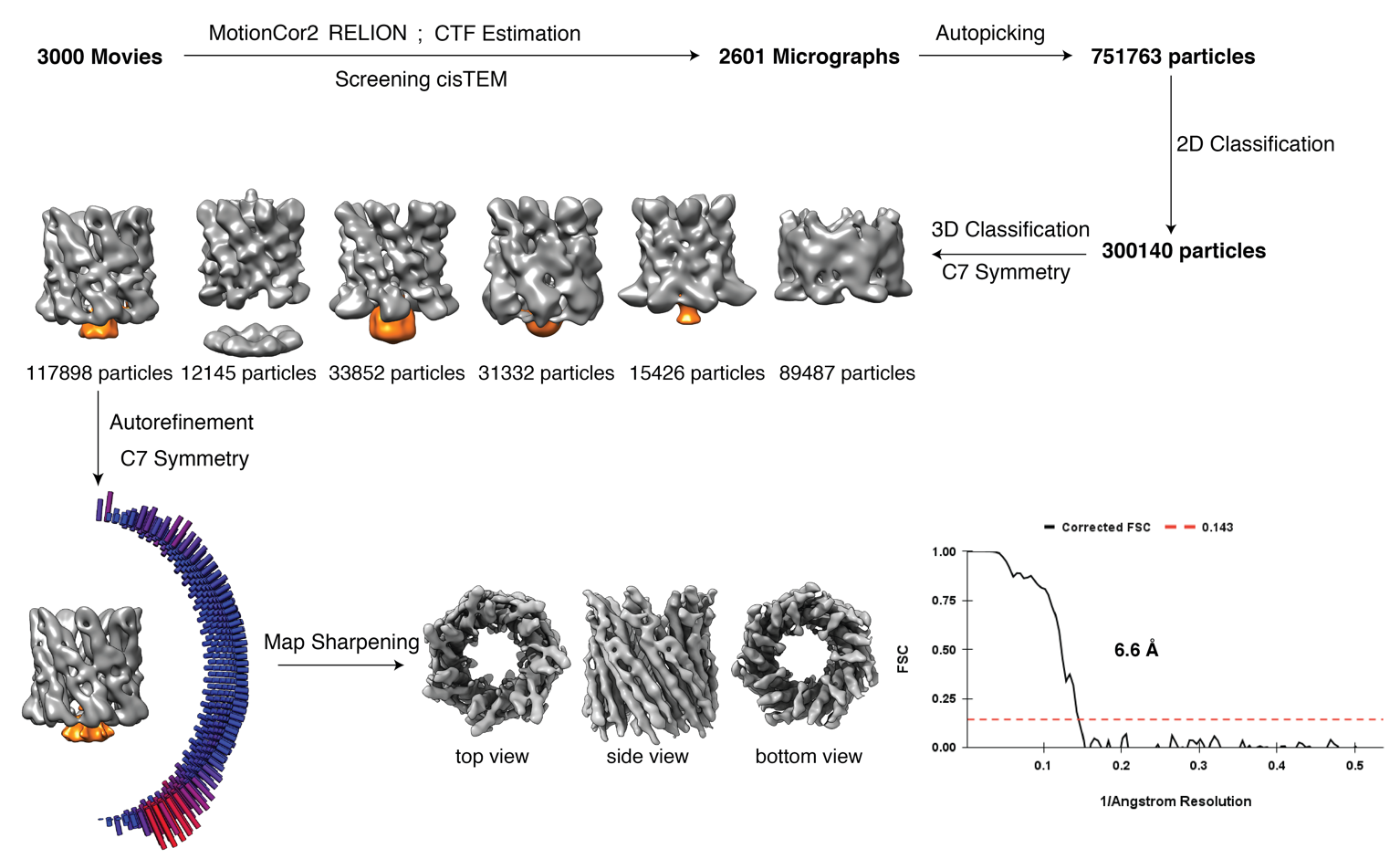

**Supplementary Figure 7**: Cryo-EM data processing pipeline followed for PA-EspB_1-332_ dataset. Orange color has been used to highlight the additional density obtained at the bottom of the map. Gold standard Fourier Shell Correlation (FSC) calculation shows a resolution of 6.6 Å. See Methods for extended procedures.

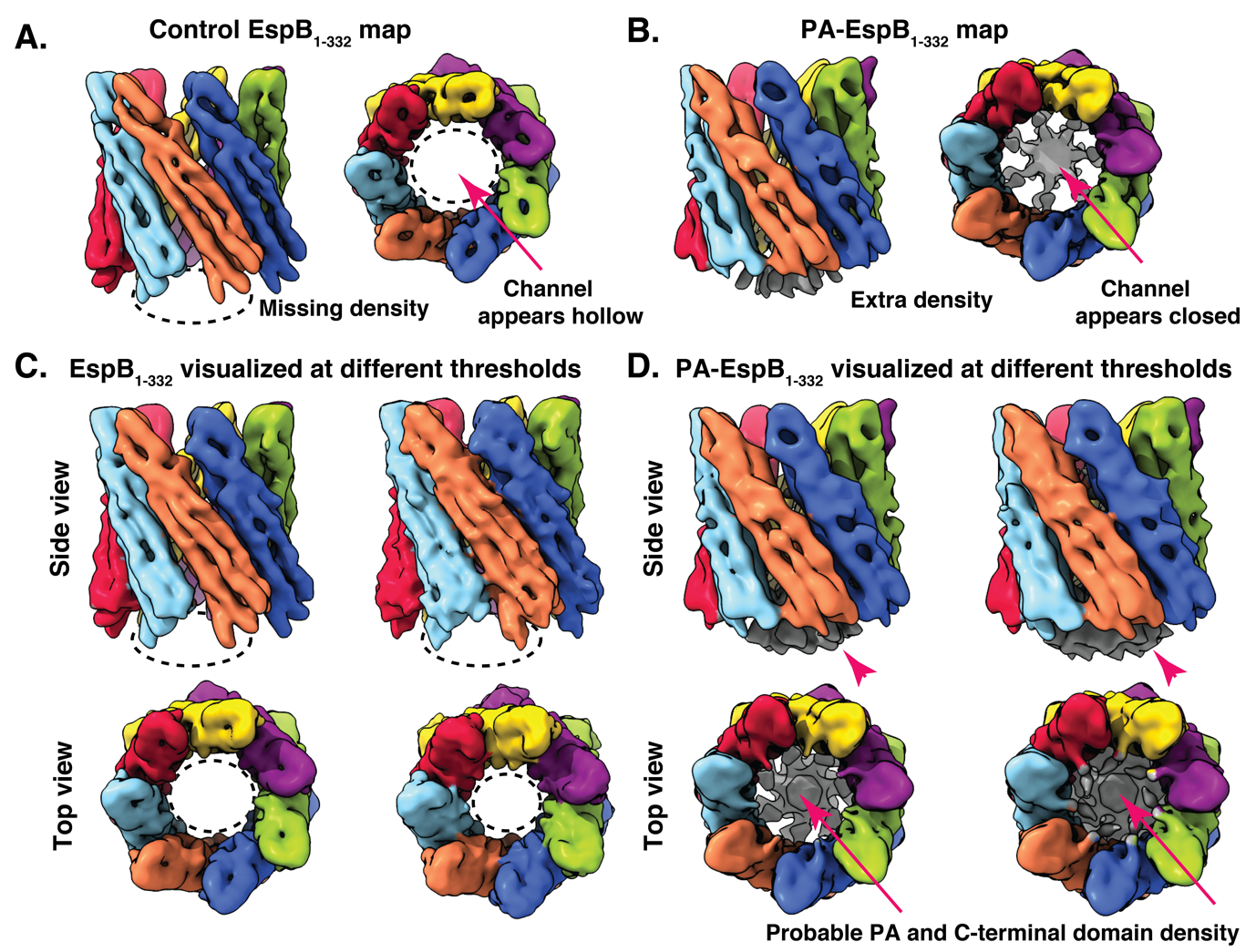

**Supplementary Figure 8:** **Comparative analysis of EspB_1-332_ structures obtained without and with PA.** (A) Side and top views of a 10 Å Gaussian low-pass filtered map of control EspB_1-332_ shows only the N-terminal domain. (B) Side and top views of a 10 Å Gaussian low-pass filtered map of PA-EspB_1-332_ shows a firm density at one end of the channel, along with the N-terminal domain. (C) Boosting the volume threshold of filtered control map does not make the additional density appear. Upper panel shows the volume boosted side views whereas the bottom panels represent the corresponding top views. (D) Boosting the volume threshold of filtered PA treated map shows an increase in the volume of the additional density. Upper panel shows the volume boosted side views whereas the bottom panels represent the corresponding top views.

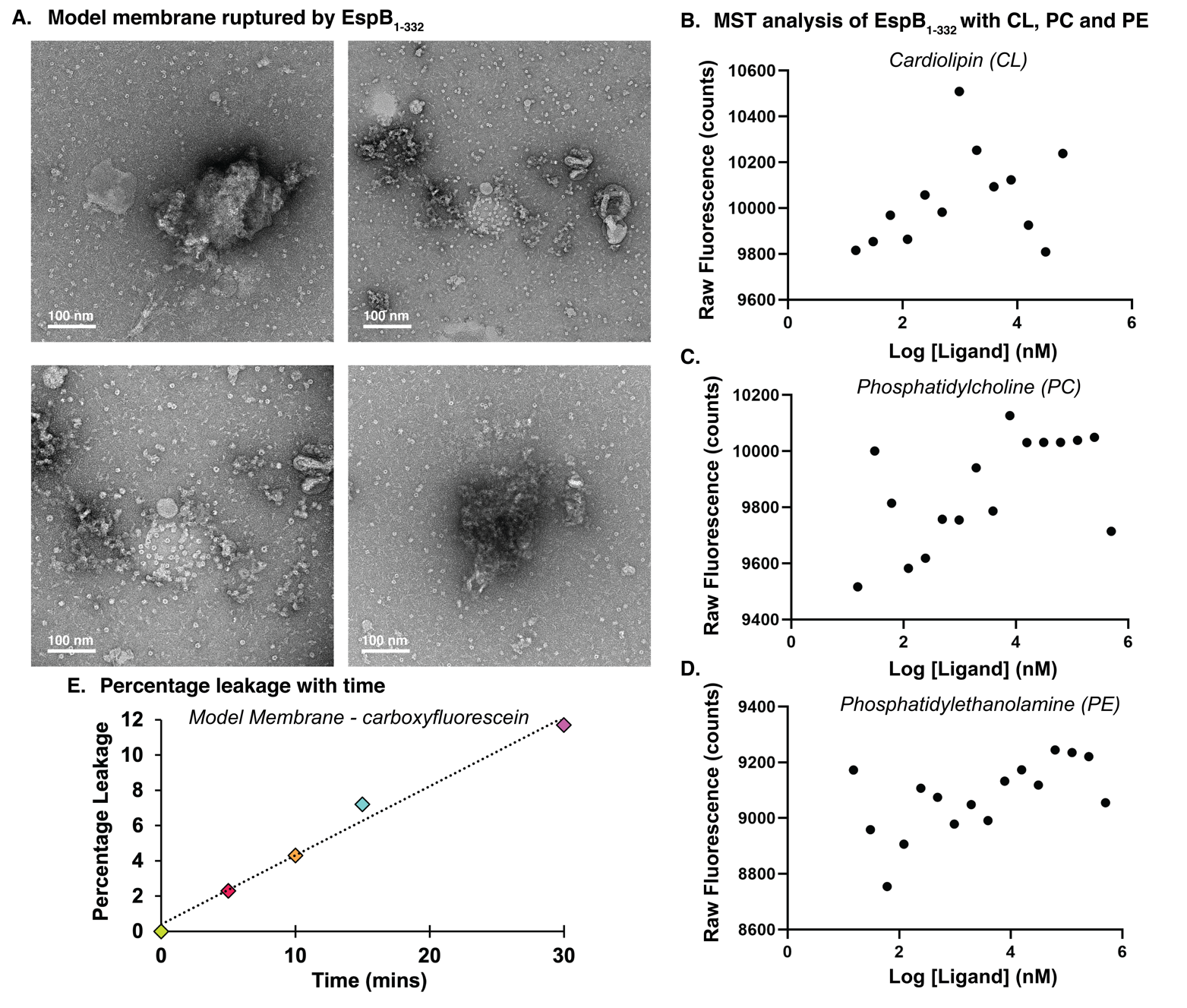

**Supplementary Figure 9**: (A) Collection of ruptured model membrane observed after incubation with EspB_1-332_. (B)-(D) Failure to observe binding affinity with cardiolipin, phosphatidylcholine and phosphatidylethanolamine, respectively. (E) Graph showing correlation between fraction of leaky liposomes and time, in a range of 0 to 30 mins.

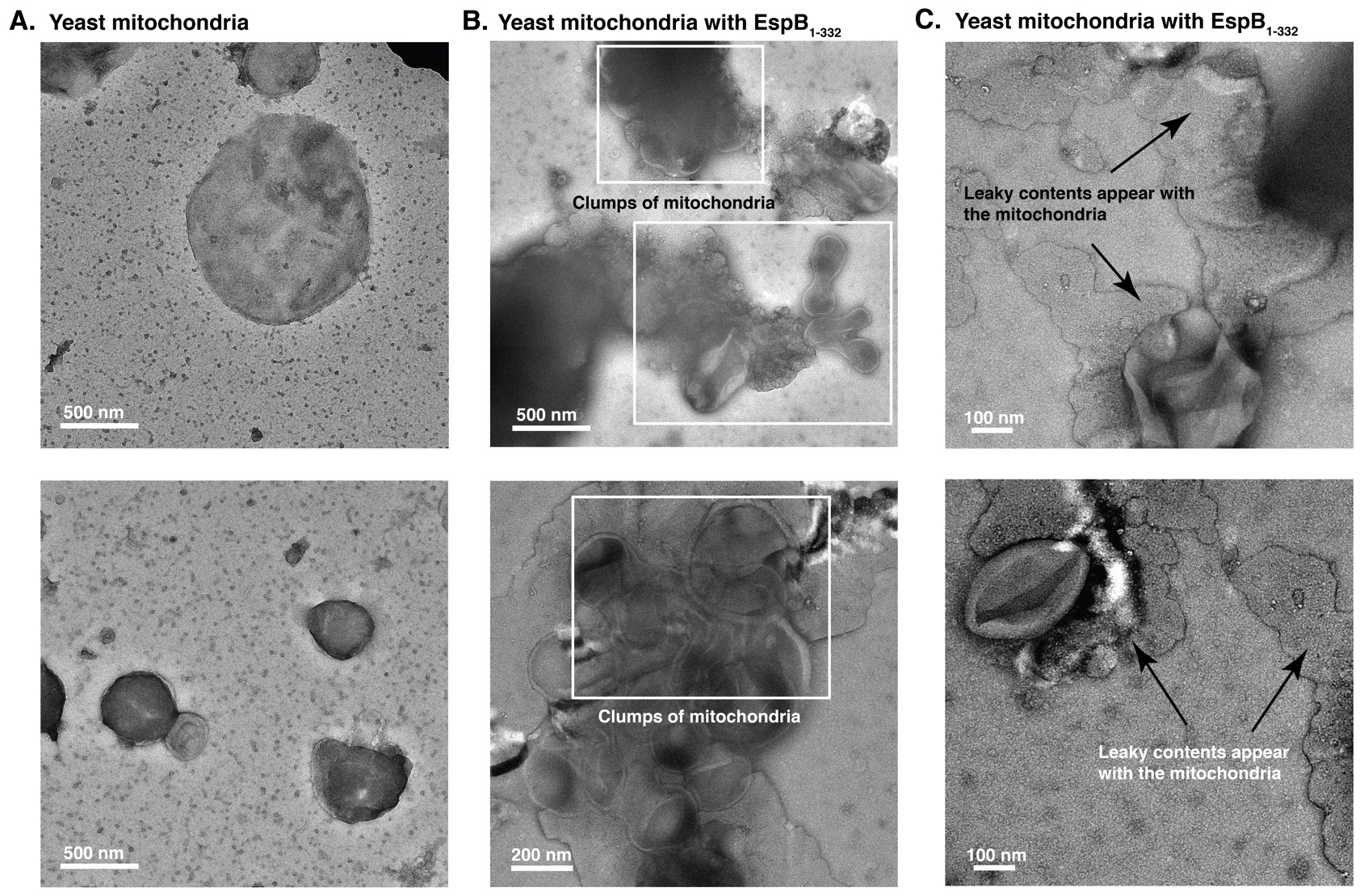

**Supplementary Figure 10**: **NS-TEM visualization of yeast mitochondria without and with EspB_1-332_.** (A) Control mitochondria observed in different areas. (B) Altered morphology of mitochondria post incubation with EspB_1-332_, indicating clumping of mitochondria. (C) shows dense patterns surrounding mitochondria giving an impression of leakage of organelle contents.
